# Efficacy of neurofeedback training for improving attentional performance in healthy adults: A systematic review and meta-analysis

**DOI:** 10.1101/2023.03.19.533384

**Authors:** Ikko Kimura, Hiroki Noyama, Ryoji Onagawa, Mitsuaki Takemi, Rieko Osu, Jun-ichiro Kawahara

## Abstract

This systematic review and meta-analysis examined the NFT effects on attentional performance in healthy adults. Six databases were searched until June 2022 to identify parallel randomized controlled trials (RCTs) evaluating attentional improvements after NFT. Risk of bias was assessed using the Cochrane Collaboration tool. We identified 41 RCTs for qualitative synthesis and 15 RCTs (569 participants) for meta-analysis. The overall NFT effect on attentional performance was significant (standardized mean difference = 0.27, 95% confidence interval = 0.10–0.44). However, no significant pooled effect was found within the trials comparing its effect with sham NFT (8 RCTs). Additionally, variable effects were observed on individual subsets of attentional performance. Further sham-controlled RCTs are required to validate the improvement of attentional performance with NFT.

## 1 Introduction

Attention is essential for cognitive functions and complex behaviors. It helps to select contextually critical information, ignore irrelevant information, and maintain or control task-relevant goals, thoughts, and emotions. Given its importance in cognitive processes, improving attentional performance would extensively benefit in daily life processes, including enhancing work performance and safety.

Several researchers have attempted to clearly define the term “attention” (Egeth and Kahneman, 1975; Pashler and Sutherland, 1998; Wickens, 2002). In fact, there is no unequivocal definition of attention as it is a multifaceted term that represents several behavioral benefits, including improvement in object detection or identification, response facilitation, and successful inhibition. Nevertheless, previous studies have identified multiple subtypes of attention. For instance, Posner and colleagues defined three distinct subtypes of attention: executive function, spatial orientation, and arousal (Posner and Petersen, 1990). Executive function establishes and maintains task-relevant goals while preventing interferences and conflicts. Spatial orientation localizes a target and prioritizes processing toward that target, irrespective of foveal fixation if the target is within the visual field. Arousal sustains the optimal level of alertness to effectively perceive high-priority signals and prepare an appropriate response. Posner and colleagues also argued that these three subtypes are mediated by largely independent brain networks (Fan et al., 2002; Posner and Petersen, 1990).

It is noteworthy that multiple studies have uncovered new aspects of attention since Posner and colleagues’ proposal. Several researchers argue that spatial orientation consists of two or three components (Corbetta et al., 2008; Luck et al., 2021; Summerfield and Egner, 2009). Others maintain that there is a substantial overlap between attention and executive functions (Diamond, 2013; Miyake et al., 2000; Oberauer, 2019) or that attention interacts with consciousness and subjective experience (Carrasco, 2018; Chun et al., 2011). Nonetheless, the three major subsets identified by Posner and colleagues (Posner and Petersen, 1990) remain unaffected by these studies and serve as a general guide to interpret the effect of attention on human performance (Petersen and Posner, 2012).

Neurofeedback training (NFT) is used to intervene in the activity of a specific brain region or network and hence is a viable way to improve attentional performance. NFT trains the participants or patients to self-regulate a specific brain activity. This can be achieved by monitoring each individual’s brain activity in real time with a device (i.e., electroencephalography [EEG], functional magnetic resonance imaging [fMRI], or near-infrared spectroscopy [NIRS]), and then by providing feedback with visual or (and) auditory stimuli. Since the first report showing the potential of EEG-based NFT to modulate the brain activity in humans (Kamiya, 1968), multiple NFT protocols have been developed to enhance attentional performance in healthy adults (See Gruzelier, (2014) for a review).

NFT targeting to increase sensorimotor rhythm (SMR) power is a major protocol used to increase attentional performance. SMR is a low-end beta band (typically 12– 15 Hz) derived from sensorimotor cortex. This rhythm was shown to occur immediately prior to the performance of certain goal-directed behaviors (Wyrwicka and Sterman, 1968). After the SMR protocol of NFT was shown to reduce inattention and hyperactivity in patients with attention-deficit/hyperactivity disorder (ADHD) (Lubar and Shouse, 1976), this has been used to increase arousal (Egner and Gruzelier, 2001, 2004a). Moreover, this method was shown to increase the performance of goal-directed complex behaviors (Cheng et al., 2015; Egner and Gruzelier, 2003). NFT to increase low beta (beta1; 15–18 Hz) was also reported to increase attentional performance but different attentional metrics than NFT to increase SMR power (Egner and Gruzelier, 2004a). This suggests that different NFT protocols might enhance different components of attentional performance.

NFTs intended to increase alpha power can also improve attentional performance. Although alpha band (typically 8–12 Hz) was originally reported to be enhanced in the relaxed and eyes-closed state rather than in the alert state (Berger, 1929), this band has been also shown to be important for maintaining task performance (Hord et al., 1976; Nowlis and Kamiya, 1970). Since then, this band has been thought to be crucial for inhibiting irrelevant stimuli (top-down attention), thereby enhancing executive function or spatial orientation (Hanslmayr et al., 2005). In addition to the bands described above, several reports have also elucidated the relationship between attentional performance and frontal theta power (Cavanagh et al., 2012; Wang and Hsieh, 2013). All these protocols can be potentially used to alter attentional performance.

Recently, two meta-analyses have shown that NFT, including protocols described above, can mitigate inattention and hyperactivity in patients with ADHD (Lambez et al., 2020; Van Doren et al., 2019). However, it remains unclear whether NFT can further enhance attentional performance in healthy adults rather than alleviate the attentional deficits in clinical populations. The results of studies in patients with ADHD cannot be directly applicable to the healthy population given that psychostimulants used to treat ADHD, such as methylphenidate, may affect neural plasticity (Korchounov and Ziemann, 2011; Ridding and Ziemann, 2010). Therefore, it is essential to investigate whether NFT is an effective approach for improving attentional performance among healthy adults. This knowledge is crucial for determining the applicability of NFT in improving their performances at work (Mazur et al., 2017; Ros et al., 2009) and in sports (de Brito et al., 2022).

Here, we conducted a systematic review of the effects of NFT on attentional performance among healthy adults. A meta-analysis was also performed to quantitatively evaluate the NFT effect in enhancing attentional performance and to clarify the NFT effect on each subtype of attentional performance based on the definition proposed by Posner and colleagues, the effect with each type of NFT protocol (e.g., enhancing SMR or alpha power), and the effect over each type of control condition. The present study differs from two previous systematic reviews on the effects of NFT on attentional performance (Da Silva and De Souza, 2021) and executive function (Da Silva and De Souza, 2021; Viviani and Vallesi, 2021) in healthy adults for the following reasons. First, neither conducted a meta-analysis. Second, the effects of NFT on the individual subsets of attentional performance (executive function, spatial orientation, and arousal) and which protocol of NFT has been used to intervene on each subset were not examined. Third, the superiority of NFT over each type of control condition (e.g., sham-NFT, general training methods such as video game play (Nouchi et al., 2021) or meditation (Egner and Gruzelier, 2004b; Nouchi et al., 2021), and no intervention) has not been assessed. This is worthwhile because comparing the efficacy of NFT with sham-NFT is crucial for clarifying the effect specific to modulating the targeted brain activity (Schönenberg et al., 2017), while comparisons with general training methods are essential for demonstrating superiority over other alternatives.

## 2 Methods

This systematic review and meta-analysis were conducted in accordance with Chapter 4 of the Minds manual for clinical practice guideline development (Minds Manual Developing Committee, 2021). This study was not pre-registered.

### 2.1 Database search

Two of four authors [IK, HN, RiO, and JK] independently searched for English and Japanese articles investigating NFT effects on attentional performance using the English databases (PubMed [IK and HN], Scopus [IK and HN], Web of Science [IK and HN], and APA PsycInfo [RiO and JK]), the Japanese database for scientific studies (JDream Ⅲ [RiO and JK]), and the Japanese database for medical trials (Ichu-shi [RiO and JK]). Searches of PubMed and Web of Science were conducted in May 2022, while searches of all other databases were conducted in June 2022. Search strings included both attention-AND NFT-related terms. Attention-related terms were “attention*” OR “wakefulness” OR “executive function,” while NFT-related terms were “neurofeedback” OR “neuro feedback” OR “brain machine interface” OR “brain–computer interface” OR (“feedback” AND EEG OR electroencephalogra*). To further eliminate clinical and animal studies, we excluded if the following keywords were present in the title: (ADHD OR disorder* OR deficit* OR patient* OR rehabilitation OR epilep* OR disease* OR depress* OR injur* OR damage*) OR (animal* OR rodent* OR monkey* OR rat OR rats OR macaque*). These searches were conducted in English using PubMed, Scopus, Web of Sciences, and APA PsycInfo, whereas JDream Ⅲ and Ichu-shi were searched using equivalent Japanese terms (See Supplementary Table 1 for the exact search queries used for each database).

### 2.2 Screening procedures

We then screened the articles retrieved from all databases in two phases. First, we removed articles that did not meet eligibility criteria based on title and abstract (first phase), and then inspected the remaining full articles for eligibility (second phase). During the second phase of screening, the following data were extracted and considered for eligibility: study design, participant demographics, intervention type, control condition, and outcome measures related to attentional performance. The eligibility criteria used in both phases of screenings were as follows: (1) Populations: healthy adults aged 18–64 years; (2) Intervention: NFT using a non-invasive measure of brain activity to improve attentional performance; (3) Comparison: any type of control condition; (4) Outcome: measurements reflecting attentional performance to assess the effect of NFT; (5) Study design: parallel randomized control trial (parallel RCT); (6) Source: Published in a scholarly peer-reviewed journal and written in English or Japanese. Three authors (IK, HN, and RyO) were involved in the screening procedure. We divided the articles into three groups for each phase, and each group was screened independently by two of three authors ([IK and HN], [HN and RyO], or [RyO and IK]). In case of disagreement between these two authors, another author reviewed the article (RyO for the groups reviewed by [IK and HN], IK for those by [HN and RyO], HN for those by [RyO and IK]) to reach a consensus. If the three authors still disagreed on eligibility, the other authors inspected the articles, and all six authors reached agreement through further discussion. Of the eligible studies, we excluded from the subsequent meta-analysis if (1) data were not available to calculate the effect size between experimental and control conditions or (2) the study was judged to have a high risk of bias (See Section 2.4. for detailed definitions of risk of bias).

### 2.3 Data extraction

Two authors (IK and HN) extracted the data from each eligible article for qualitative synthesis and meta-analysis. For qualitative synthesis, the following details were extracted: participant demographics, intervention, control condition, outcome measures of attentional performance, and adverse events if any. Adverse events are any unfavorable medical events that occurred during a study period, such as headache, nausea, or dizziness (Cumpston et al., 2019). The attentional outcome measures in each study were further categorized according to the targeted attentional performance subsets as follows: executive function (e.g., Flanker effect: measuring interference and response competitions from spatially flanking irrelevant distractors toward targets (Eriksen and Eriksen, 1974; Eriksen, 1995); Simon effect: measuring interference and response competitions from spatially congruent but irrelevant distractors, (Simon and Rudell, 1967); Navon task: measuring interference and response competitions from global (or local) distractors (Navon, 2003, 1977); Psychological refractory period effect: measuring temporal separations between two processes that are assumed to share common attentional resources (Pashler, 1994; Welford, 1952); Dual-task performance: measuring interference from one task to another that is assumed to share common attentional resources (Allport et al., 1972; Logan and Gordon, 2001); Task switching: measuring a temporal delay reflecting time to reconfigure one set of attention to another (Monsell, 2003; Vandierendonck et al., 2010); Working memory span task: measuring attentional capacity of working memory during reading sentences (Daneman and Carpenter, 1980); Operation span task: measuring attentional capacity of working memory during using executive function while concurrently maintaining information temporarily (Draheim et al., 2022; Turner and Engle, 1989); Paced auditory serial attention test: measuring attentional capacity during calculating auditorily presented numbers while listening to another number (Gronwall and Sampson, 1974; Tombaugh, 2006); Trail making test: measuring executive function by assessing performances in tracking visually scattered alphabets and numbers sequentially but interchangeably (e.g., 1-A-2-B-3-C …) (Tombaugh, 2004); N-back task: measuring working memory capacity by assessing performance in searching for a target letter or number repeated in a sequence (Jaeggi et al., 2008; Kirchner, 1958); Attentional blink: measuring temporal limitation of selective encoding on a second target presented in a rapid sequence of stimuli (Dux and Marois, 2009; Raymond et al., 1992); Change detection: measuring maintenance of visuospatial array over time, (Pashler, 1988; Rensink, 2003); and Priming: measuring the effect on behavior or perception after exposure to certain stimuli (Neely, 1977)), spatial orientation (e.g., Spatial cueing: measuring spatial allocation of attentional resources (Corbetta et al., 2002; Posner, 1978); Visual search: measuring distribution and focus of attentional resources over spatial array (Neisser, 1964; Wolfe, 2020); Multiple object tracking: measuring distribution and focus and maintenance of attentional resources for moving objects over time (Pylyshyn and Storm, 1988; Scimeca and Franconeri, 2015); Subitizing: measuring parallel access to multiple spatially distributed items (Balakrishnan and Ashby, 1992; Pylyshyn, 1989); Dichotic listening test: measuring ability to selectively listen to one message from one ear while ignoring the other from the other ear (Cherry, 1953)), and arousal (e.g., Sustained attention to a response task: measuring task maintenance and response inhibition by requiring repetitive responses to nontarget stimuli except for one critical target (Robertson et al., 1997); Continuous performance task: measuring sustained attention by requiring responses only to targets while refraining from responding to nontargets (Braver, 2012; Rosenberg et al., 2013); and Mind wandering: measuring state of mind whether it remains on the current task or shifts away (Schooler et al., 2004; Smallwood and Schooler, 2015)). Two authors (IK and HN) independently inspected the measured subsets. If judgements were inconsistent, four authors (IK, HN, RiO, and JK) discussed the classification until an agreement was reached.

Outcome measures of attentional performance after the last NFT session were further extracted for the meta-analysis. Here, as outcome measures, we regarded the change in scores from before NFT, and, if such data were unavailable and the scores before NFT were not significantly different between the interventional and control groups, the scores after NFT. We regarded every experiment in each paper as a single trial, form which the following values were extracted from each for the meta-analysis: mean and standard deviation of the attentional outcome measures after the last NFT session and the number of participants in the experimental and control groups. From either these values or t-values, the standardized mean difference (SMD) was calculated using Hedges’ g. If neither SMDs nor t-values were available, we contacted the corresponding author of the study for data request. For consistency across studies, we reversed the sign of SMD if a lower value reflected higher attentional performance (e.g., reaction time, tiredness, or sleepiness), so that higher SMD consistently indicates that NFT increased attentional performance compared to controls.

### 2.4 Risk of bias assessment

Two authors (IK and HN) assessed the risk of bias using the Cochrane Collaboration’s tool (Higgins et al., 2011) towards all the studies whose SMDs are available. This tool classifies risk of bias as high, unclear, or low in six domains: selection bias, performance bias, detection bias, attrition bias, reporting bias, and others (including conflict of interest, early trial termination, and incorrect sample size determination or statistical methods) (For the exact criteria for determining high or low in each domain, see Supplementary Table 2). Following Minds Manual Developing Committee, (2021), each domain was scored two points for high, one for unclear, or zero for low, and then points were summed to yield an overall risk score. We then classified overall risk as high if the score was more than seven or more than two domains were high, as low if total score was less than five and less than two domains were high, or as moderate in other cases. Individual studies judged to own an overall low or moderate risk of bias were included in the meta-analyses. A summary table on risk of bias evaluation was generated using *robvis* (https://mcguinlu.shinyapps.io/robvis/; McGuinness and Higgins, 2021).

### 2.5 Statistics

The basic characteristics of studies that passed the second screening were tabulated, including participant demographics, adverse events, targeted brain location and activity for NFT, type of control condition, and assessed subset of attentional performance (executive function, spatial orientation, and/or arousal) (Table 1). We then performed a standard pairwise meta-analysis using a random-effects model. All results for the same outcome and population were pooled into a single dataset. Publication bias was visualized using a funnel plot and evaluated using Begg’s and Egger’s tests (with *P* < 0.1 on both tests considered significant following the criterion of Chapter 4 of the Minds manual for clinical practice guideline development (Minds Manual Developing Committee, 2021)). Heterogeneity across studies was assessed using Cochran’s Q and the I^2^ statistic.

**Table 1.**
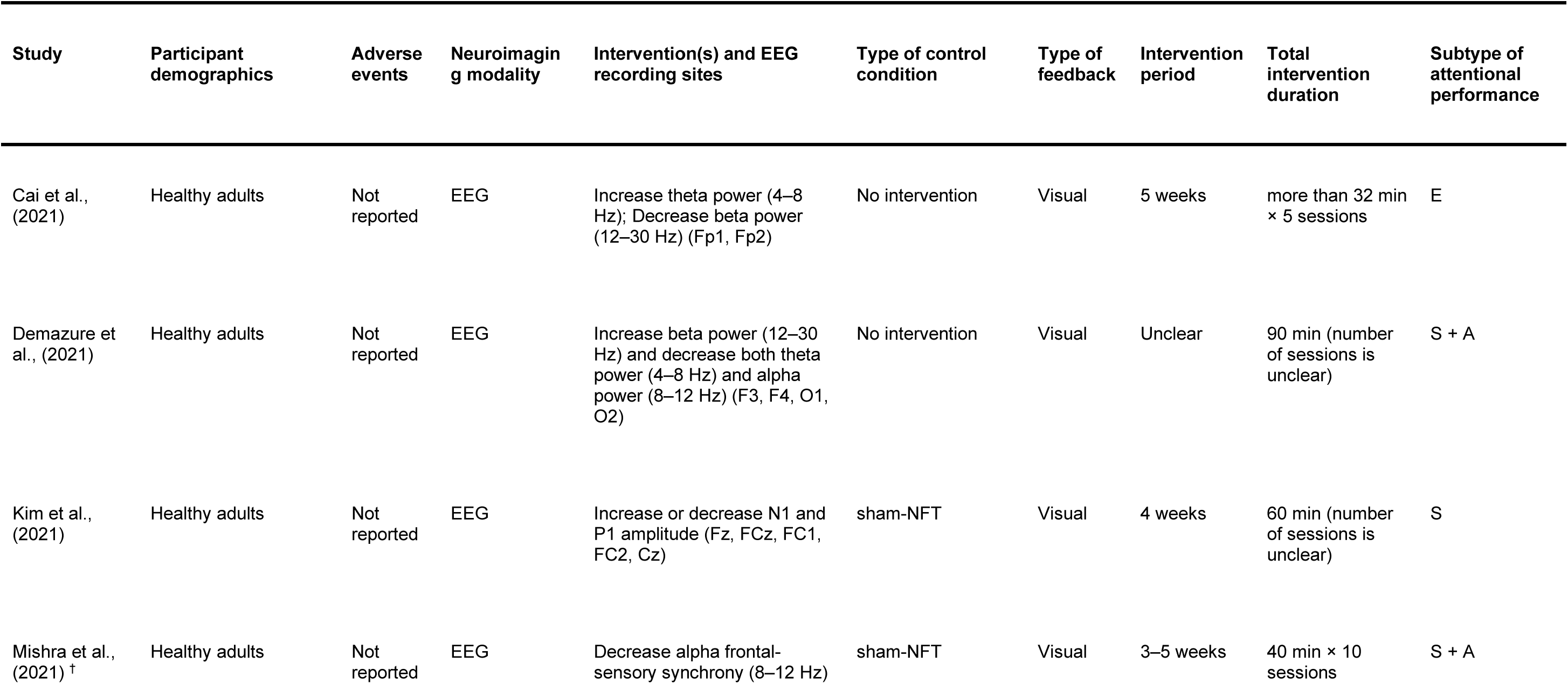
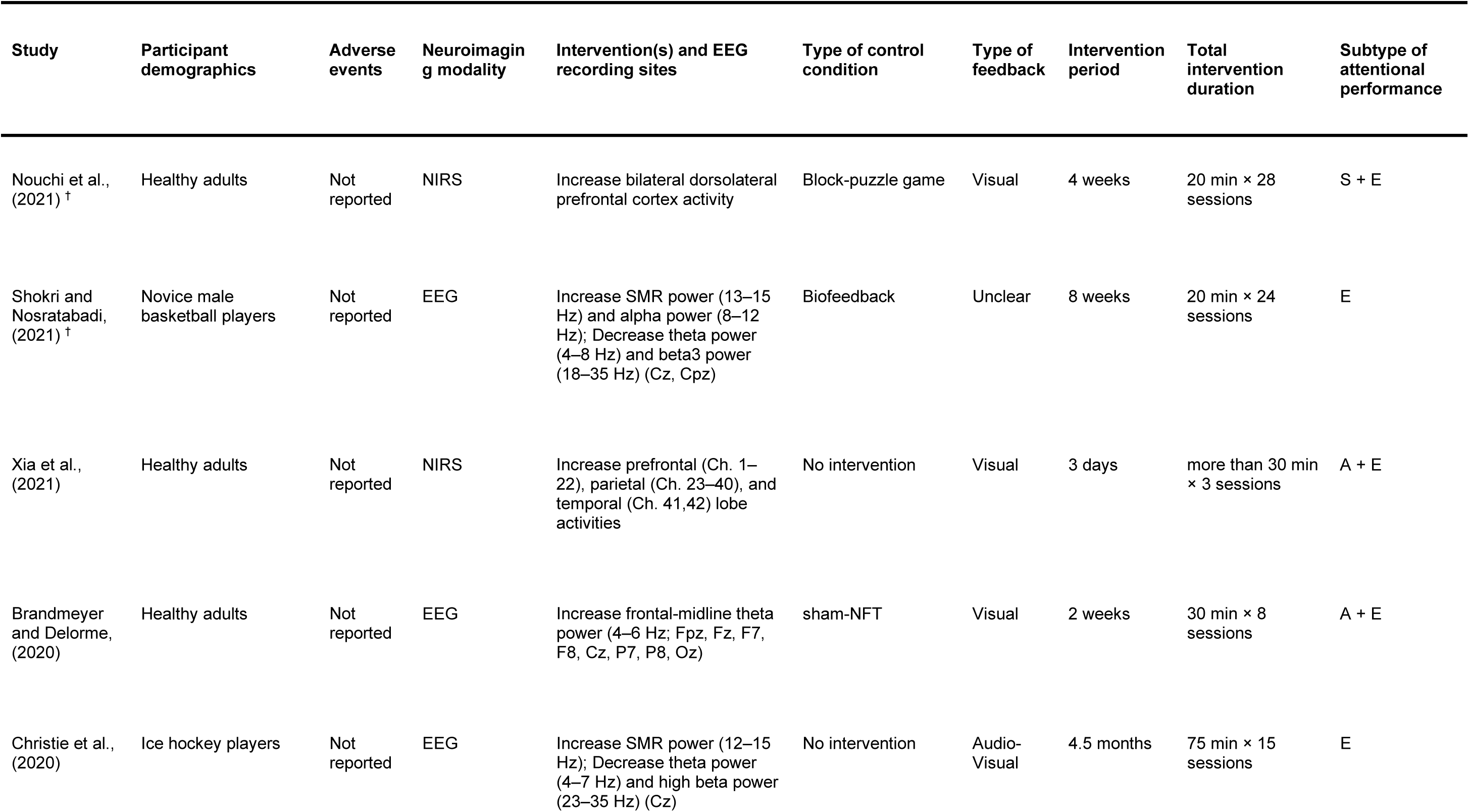
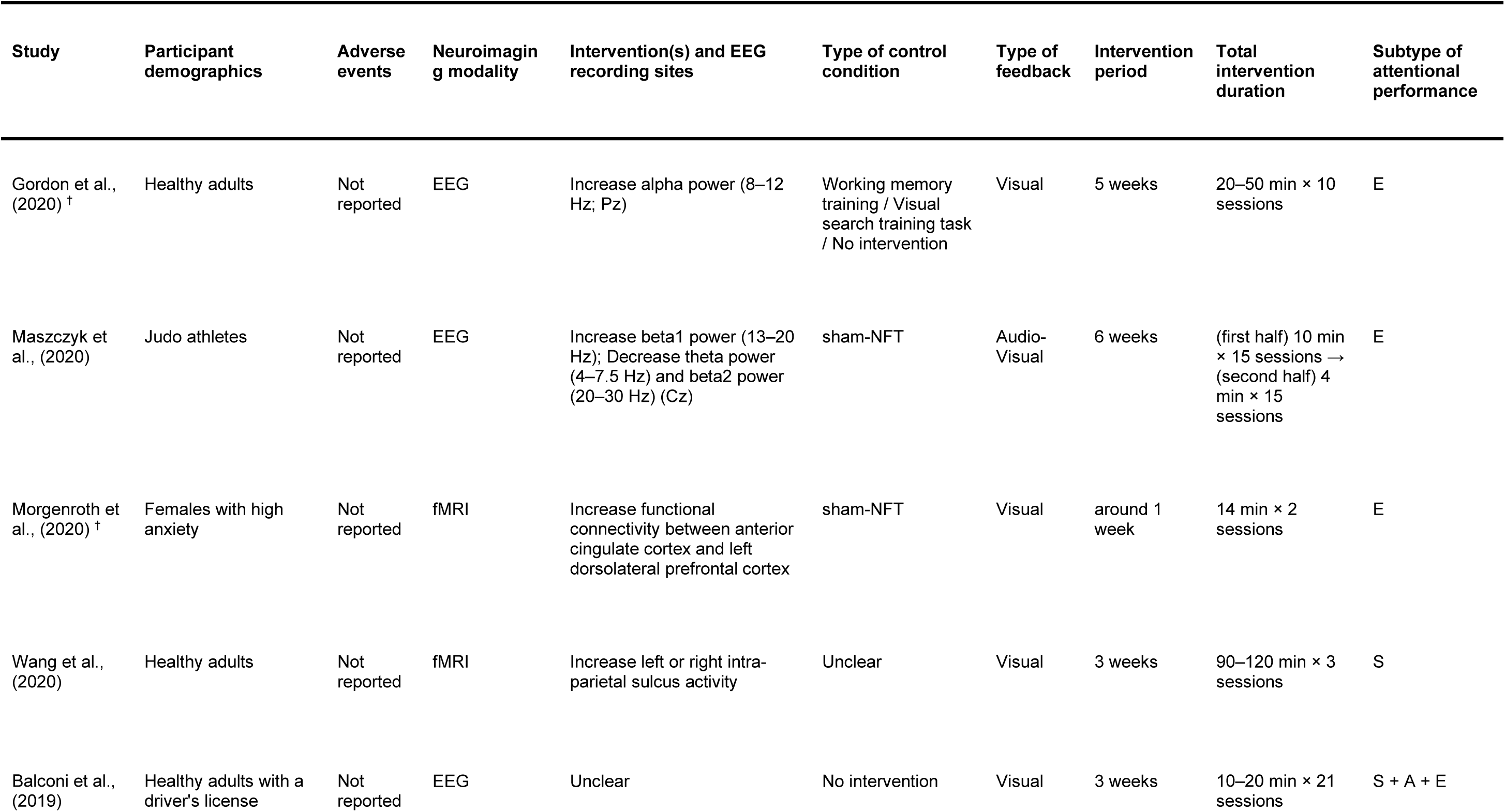
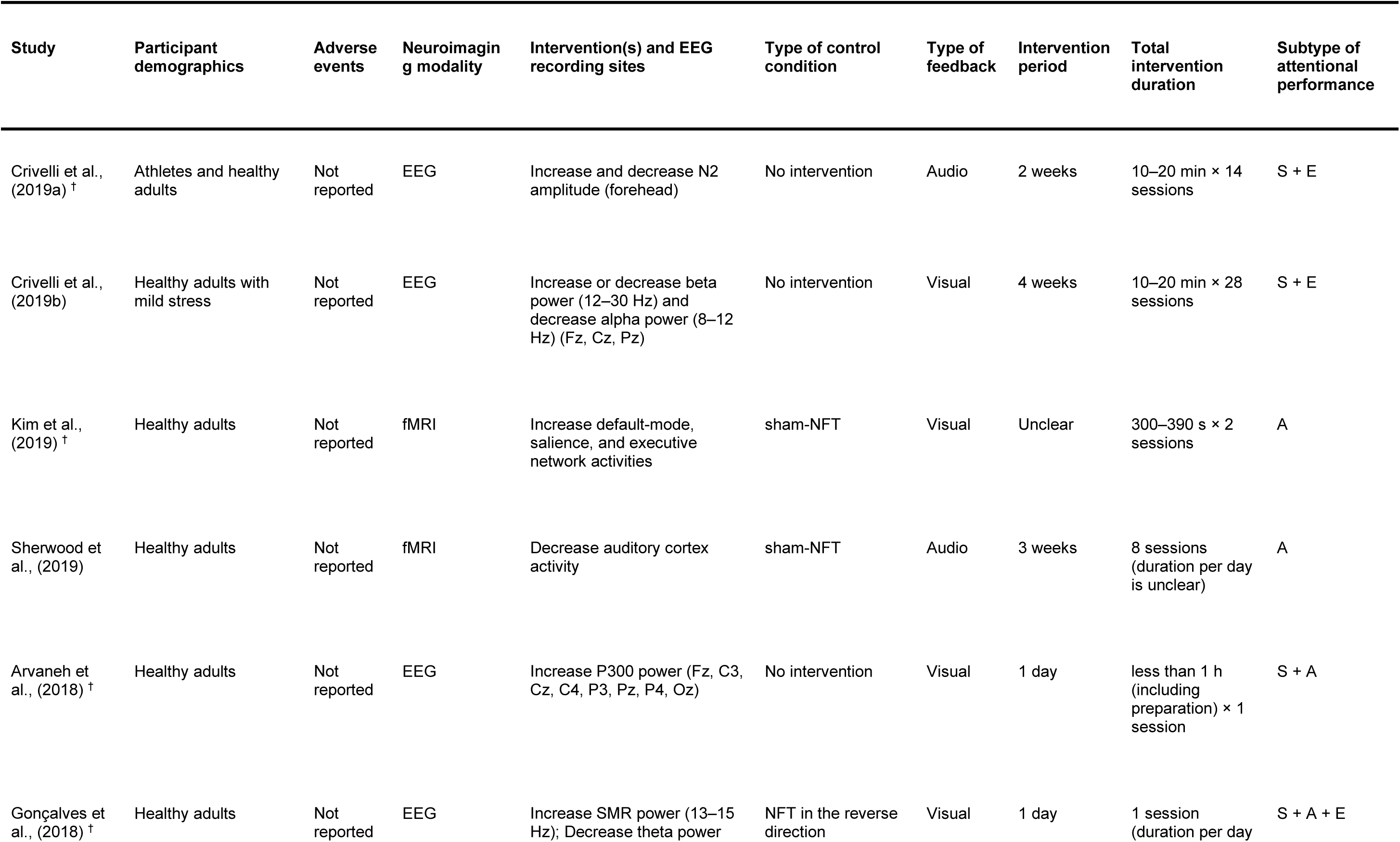
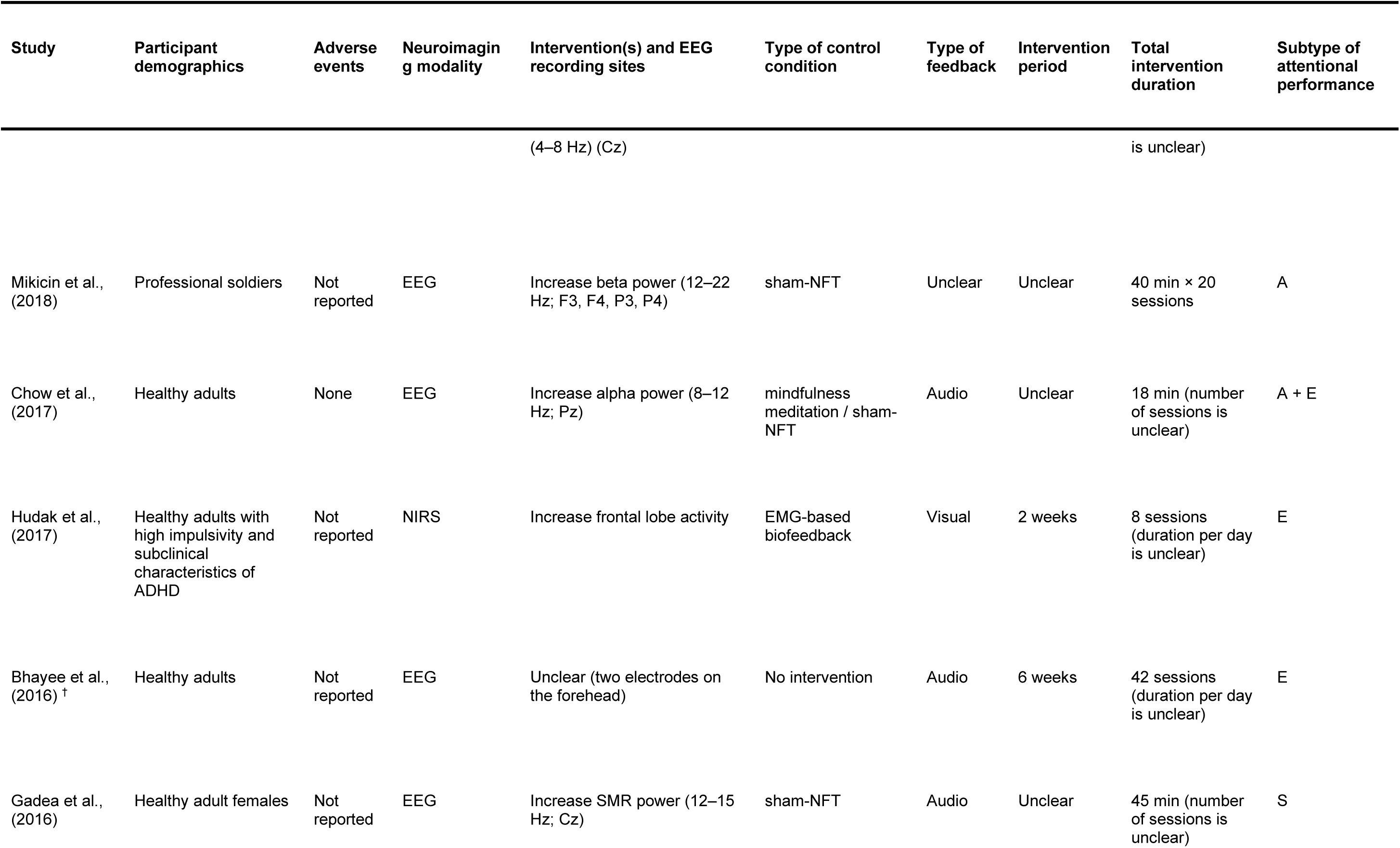
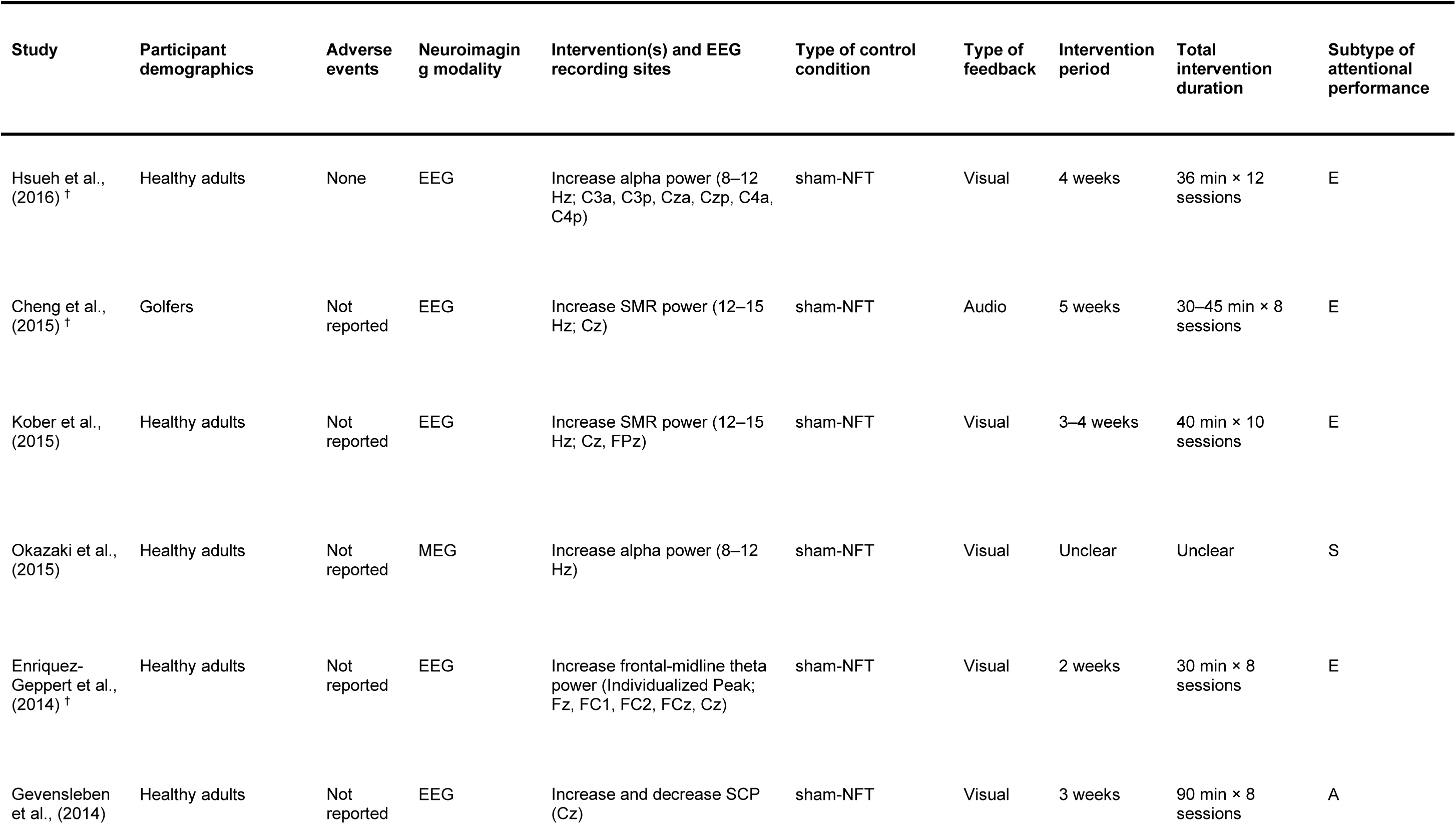
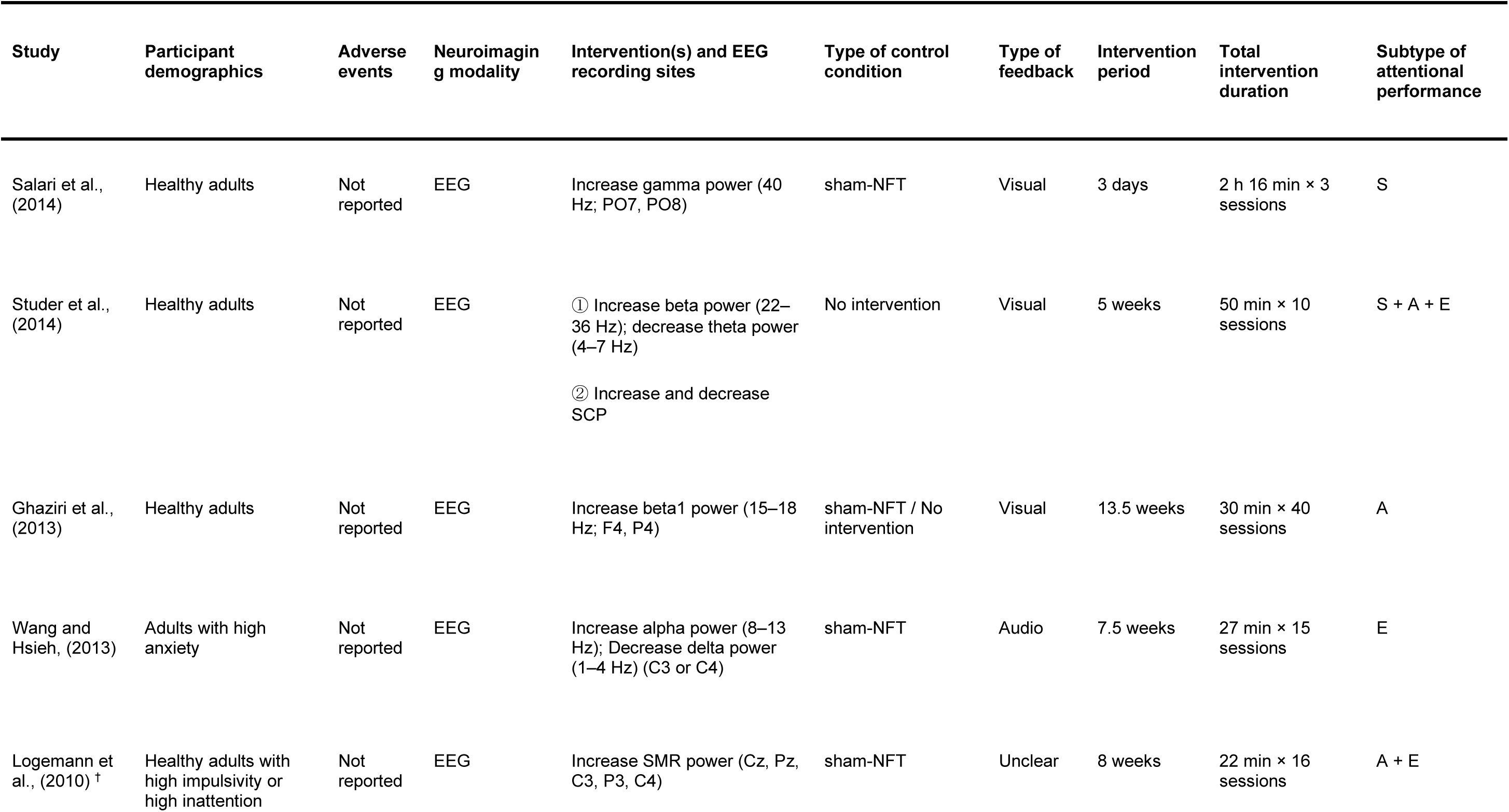
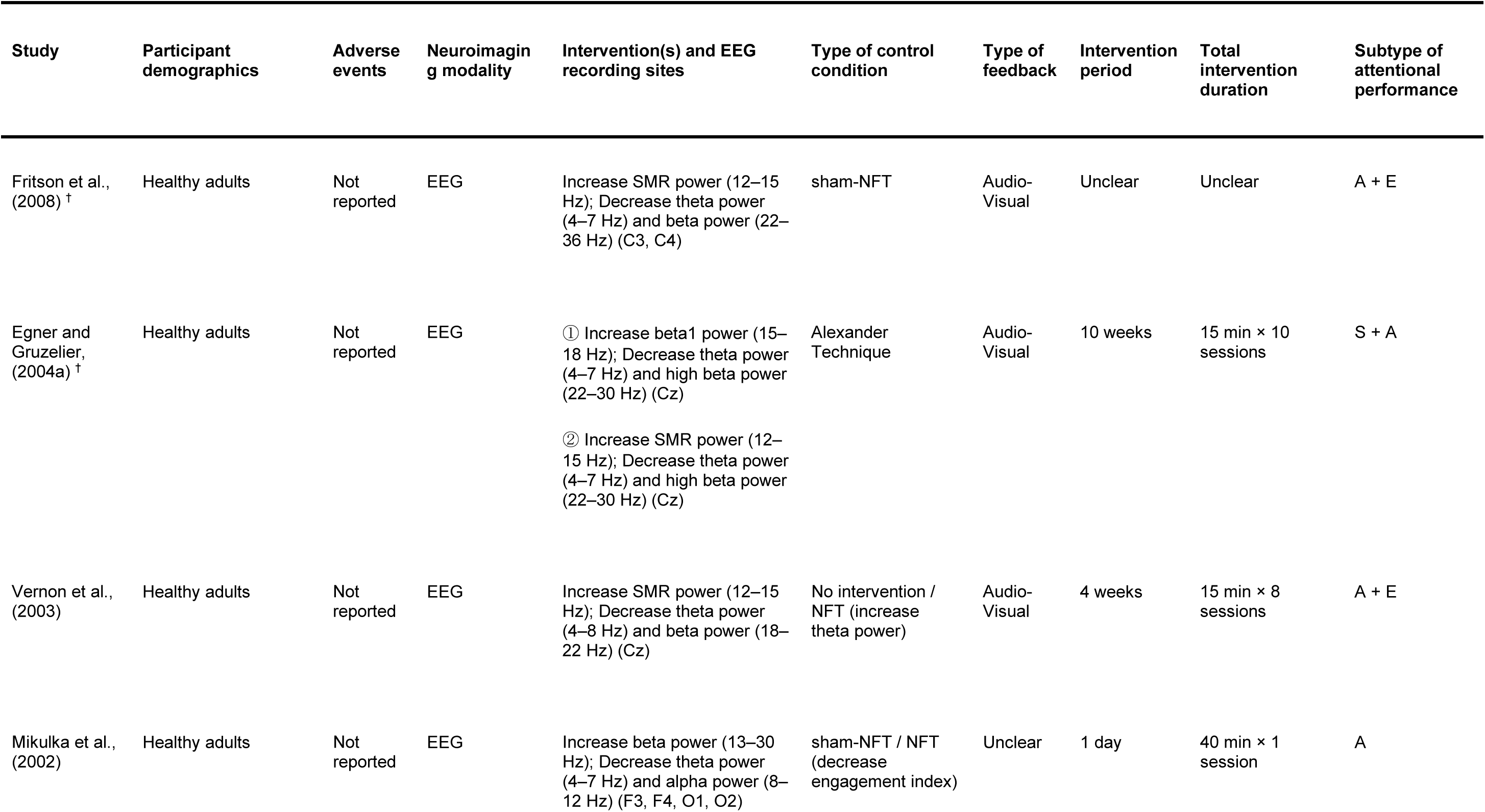
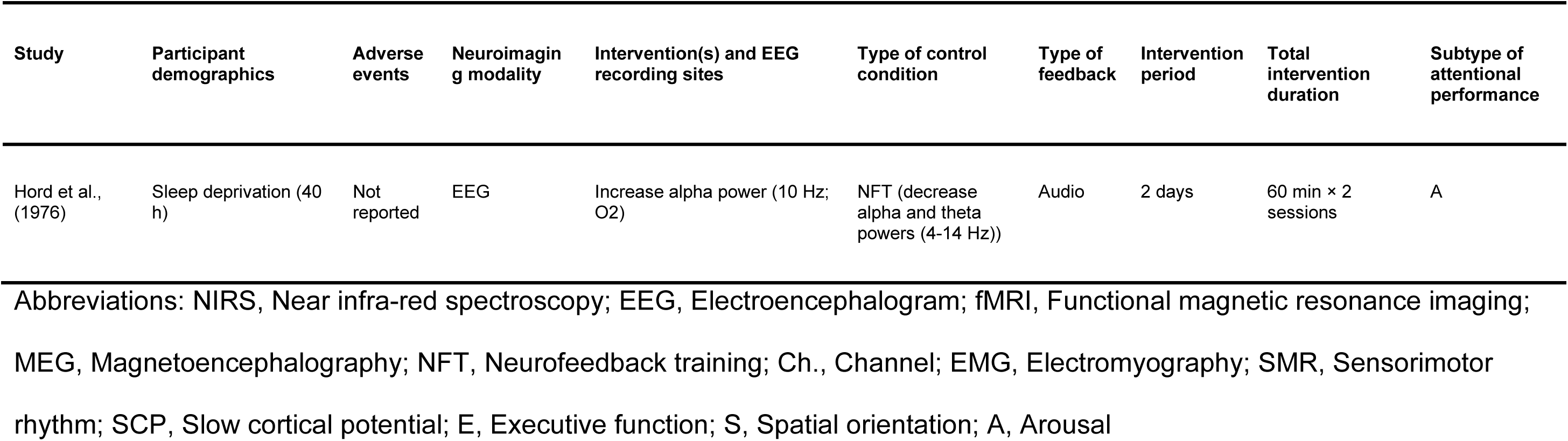
Experimental characteristics of the studies included in the qualitative synthesis. A qualitative summary was provided for each study according to the top-line items. ^†^ Studies included in the meta-analysis.

To confirm the robustness of the result on meta-analysis, sensitivity analyses were performed. Here, we confirmed whether the result of the significance test was the same and whether the magnitude of the estimated SMD was not severely changed by the following conditions: (1) changing the analysis method to a fixed-effects model, (2) including the studies with a high risk of bias, (3) excluding the SMDs calculated from the scores after the last session of NFT instead of the change in scores, (4) restricting the studies to those performing NFT with EEG. The last one was conducted because the neural mechanisms induced by NFT may be different between NFT with EEG and those with fMRI or NIRS (Nicholson et al., 2020).

To explore dose-response relationship, a meta-regression analysis was applied to examine the correlation between the effect size and total duration (or number of sessions) of NFT in each study. Here, we defined the total duration of NFT as the duration of each session × the number of sessions. If the duration of NFT in each session was described as a range (e.g., 30–40 min), the mid-value was used (e.g., 35 min). Since the effect size may decrease with publication year (Yang et al., 2023), a meta-regression analysis was also performed to investigate the relationship between publication year and effect size.

We exploratory conducted additional subgroup meta-analyses to clarify the efficacy of NFT for each subset of attentional performance (executive function, spatial orientation, or arousal), type of outcome measure (subjective [i.e., a self-assessment through a questionnaire] or objective [i.e., a quantifiable measurement that can be recorded by a device]), type of brain activity targeted for intervention (SMR or alpha power), and type of control (sham-NFT, general training, or no intervention). Because several studies assessed multiple effects with same participant groups, towards all meta- and meta-regression analyses, we applied a cluster-robust variance estimation method (Fisher and Tipton, 2015) with the *clubSandwich* package. This method can handle multiple interdependent effect sizes within each study.

To validate whether the number of trials in each main and subgroup meta-analysis was sufficient to reliably detect the effect, we performed the power-analysis for each of these analyses. The detailed method was described in (Onagawa et al., 2023), but, in brief, the statistical power and required number of sample size were estimated from the SMD and heterogeneity (*τ*^2^) calculated from each analysis using the *dmetar* package. Alpha level was set to 0.05, and we considered statistical power (1-*β*) greater than 0.8 as sufficient to detect the effect reliably. All statistical analyses were performed using *metafor* ver. 3.4.0 (Viechtbauer, 2010), *meta* ver. 5.5.0 (Balduzzi et al., 2019), *clubSandwich* ver. 0.5.8, and *dmetar* ver 0.1.0 in R ver. 4.1.2 (https://www.r-project.org/).

## 3 Results

### 3.1 Search results

The literature review process is illustrated in Figure 1. We retrieved 3,337 articles from six databases using the indicated search strings, of which 3,159 were excluded based on the title or abstract (e.g., a case report (Mroczkowska et al., 2014), a review (Sitaram et al., 2017), an observational study (Santamaría-Vázquez et al., 2022), an interventional study but not with neurofeedback (Falcone et al., 2018)), or conference proceedings (Zhang et al., 2019)). The remaining 178 articles were then screened for eligibility by full-text assessment, of which 137 were excluded for the following reasons: study of a clinical population (1 article), age of subjects (17 articles), article type (27 articles), no NFT (10 articles), no control conditions (22 articles), no outcome measure for attentional performance (38 articles), and non-parallel RCT (22 articles). The remaining 41 articles were retained for qualitative synthesis and 15 were further included in meta-analyses. Reasons for exclusion from the meta-analyses were insufficient data reported to calculate effect sizes (24 articles) and high overall risk of bias (2 articles).

**Figure 1.**
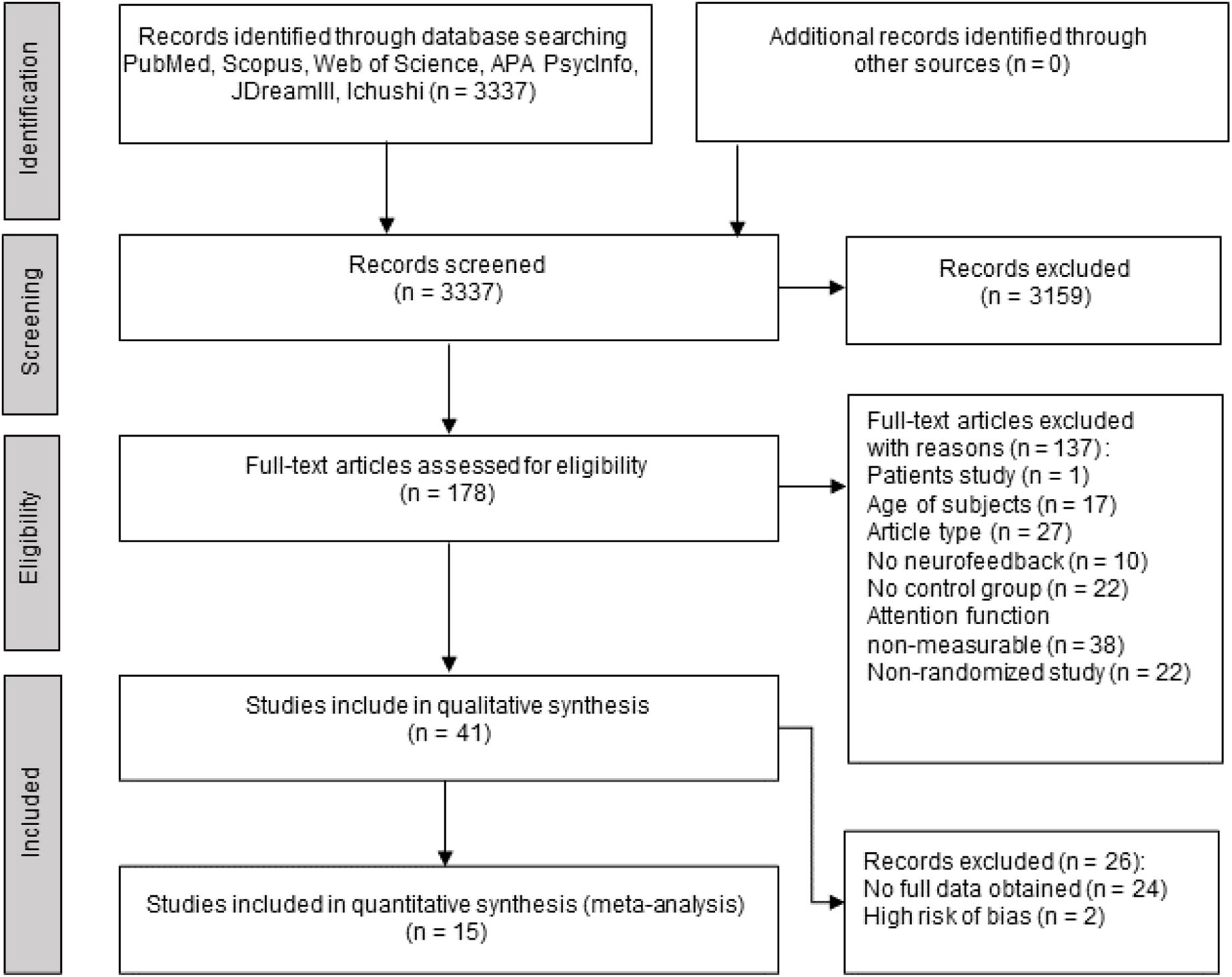
PRISMA flowchart of article search and inclusion.

### 3.2 Study and sample characteristics

Table 1 summarizes the main characteristics of all studies included in the qualitative synthesis. All were published between 1976 and 2021 (Figure 2A), with more than half appearing after 2013. Thirteen studies examined executive function alone, five examined spatial orientation alone, and seven examined arousal alone, while three examined both executive function and spatial orientation, six examined both executive function and arousal, four examined both spatial orientation and arousal, and three examined all three subsets of attentional performance (Figure 2B).

**Figure 2.**
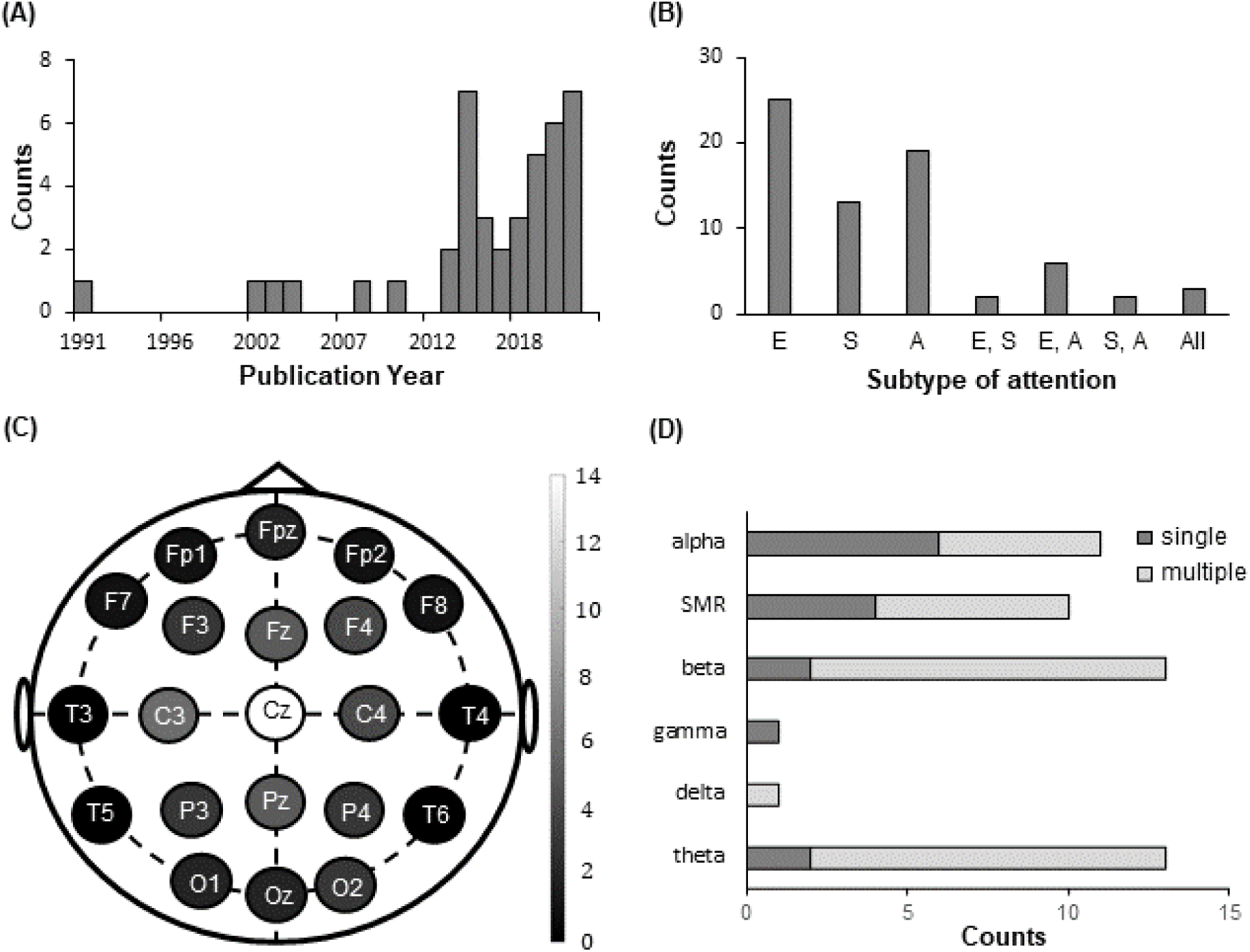
Qualitative assessment of NFT studies included in the qualitative synthesis. (A) Histogram of publication year. (B) Subset(s) of attentional performance assessed in NFT studies. (C) Channels recorded for EEG-based NFT. (D) Frequency band(s) targeted to modulate in EEG-based NFT studies. Single and multiple indicate that a band was modulated by intervening single and multiple band(s), respectively. Abbreviations: E, Executive function; S, Spatial orientation; A, Arousal; SMR, sensorimotor rhythm.

To measure the brain activity, 34 studies used EEG, 4 used fMRI, 2 used near-infrared spectroscopy (NIRS), and 1 used magnetoencephalography (MEG). More than half of the studies provided feedback with visual stimuli (24 studies), followed by audio stimuli (8 studies), and visuo-audio stimuli (5 studies). The stimulus type of feedback was unclear in four studies.

Among studies of EEG-based NFT, the vertex (Cz) was the most frequently applied electrode location, followed by the right frontal (F4), midline frontal (Fz), central (C3, C4), and midline parietal (Pz) locations (Figure 2C). The band power(s) targeted to modulate in EEG- or MEG-based NFT were alpha, SMR, beta, gamma, delta, and/or theta (Figure 2D). Most studies targeted to increase alpha power with a single band (five out of six studies), while beta and SMR powers were more often targeted to increase with multiple bands (beta power: five out of seven studies; SMR power: six out of 10 studies) than with single band. Two studies targeted to modulate slow cortical potential (SCP). To investigate the NFT effect on executive function, five, four, eight, and three studies applied NFT targeted to increase alpha, beta, SMR, and theta power, respectively, while, to explore the NFT effect on spatial orientation, two, four, three, and one studies applied NFT to increase alpha, beta, SMR, and gamma power, respectively. To evaluate the NFT effect towards arousal, three studies targeted to increase alpha, while six, five, and one studies targeted to increase beta, SMR, and theta power, respectively.

Twenty-six studies compared NFT with sham-NFT, 12 studies with no intervention, 4 studies with general training, and 2 with biofeedback without EEG. The control condition was unknown in one study.

### 3.3 Adverse events

Only two of 41 studies reported the presence or absence of adverse events, and neither reported any such events during or after NFT.

### 3.4 Risk of bias assessment

Of the 17 parallel RCTs evaluated, overall risk of bias was high in 2 studies, moderate in 13 studies, and low in 2 studies (“Overall” column in Figure 3). Only three studies explicitly described the random sequence generation method and were evaluated as low risk for selection bias (column D1 in Figure 3), while the others did not describe the allocation method and were judged as unclear risk. Only one study described the allocation concealment, which was judged as low risk for selection bias (column D2 in Figure 3). Two studies were single-blinded or unblinded and thus judged as high risk of performance bias (column D3 in Figure 3), five were double-blinded and deemed low risk, and ten did not describe the blinding procedure and were deemed as unclear risk of performance bias. Most studies did not explicitly mention the blinding status of the evaluator, so the risk of detection bias was unclear (column D4 in Figure 3). Further, one study was explicitly unblinded. Thirteen studies were judged as low risk of attrition bias (Column D5 in Figure 3), while four studies were judged as high risk since the drop-out rate was more than 10% or intention-to-treat protocols were violated. Fifteen studies had unclear reporting bias as none performed pre-registration (column D6), while one study was judged as low risk because data were collected according to the pre-registered protocol, and one was judged as high risk because the reported items were incomplete without any explicit reason. Finally, seven studies were rated high risk of other biases due to conflicts of interest (e.g., one of the authors was the employee of or the experiment was funded by a company selling neurofeedback-related devices) (three studies), early trial termination (one study), or inadequate statistical methods (e.g., no considerations for multiple comparisons) (three studies). The other ten studies were rated unclear risk due to the lack of information any on these items. Two studies with high overall risk of bias were excluded from the subsequent meta-analysis.

**Figure 3.**
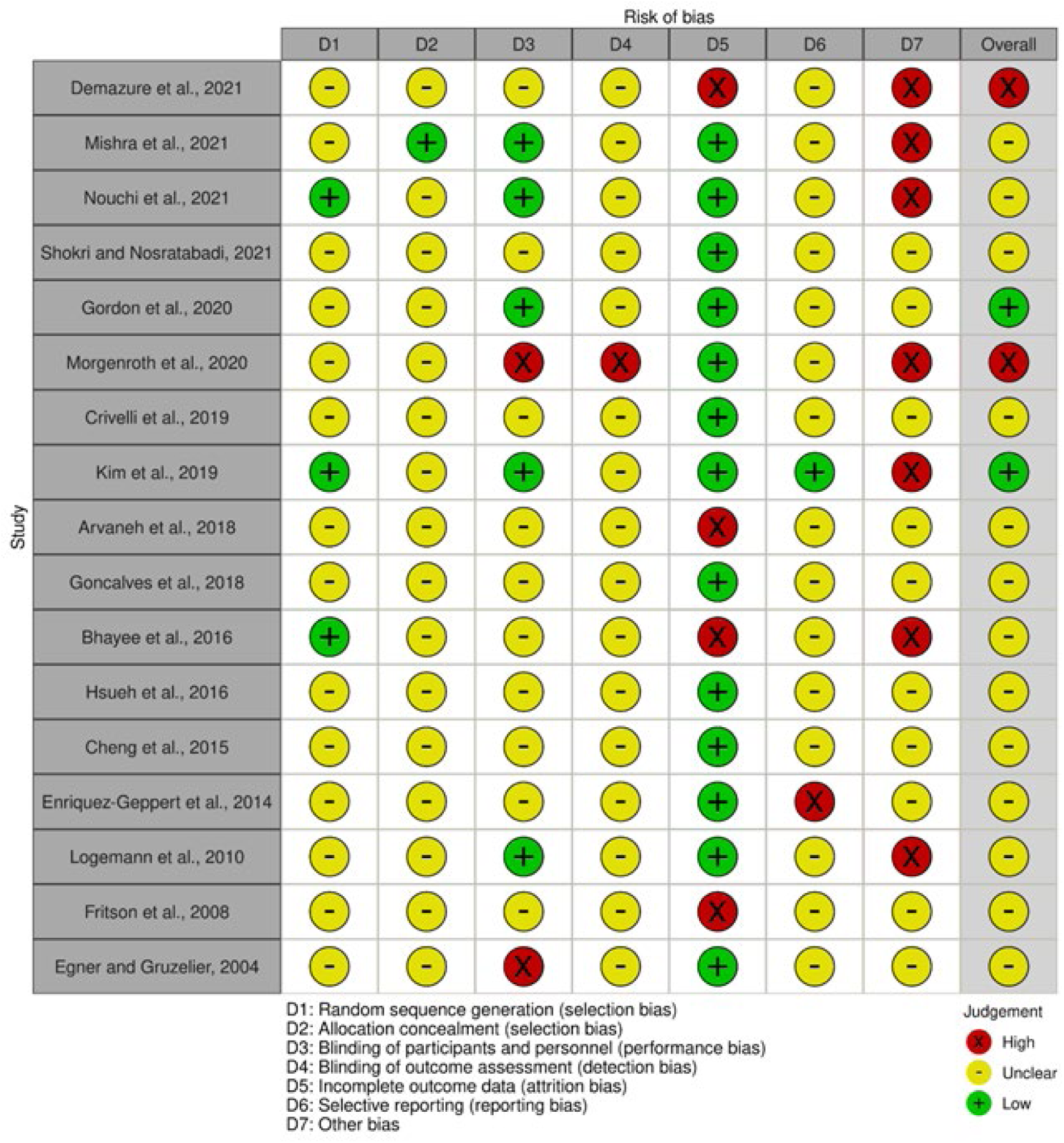
Risk of bias assessment for each study considered for the meta-analysis. The red, yellow, and green symbols in each domain column indicate high, unclear, and low risk of bias, respectively. In the overall column, red, yellow, and green indicate high, moderate, and low risk of bias, respectively. Based on this analysis, two studies (red in the overall column) were eliminated from the meta-analysis.

### 3.5 General effects of NFT on attentional performance

A total of 16 trials from 15 published studies were included in the primary meta-analysis. NFT enhanced attentional performance with small effect size (SMD = 0.27, 95% confidence interval (CI) = 0.10–0.45, *t* = 6.68, *P* = 0.0076; Figure 5A; See Supplementary Figure 1 for the forest plot). There was no significant heterogeneity across the trials (Q = 52.5, *P* = 0.93, I^2^ = 0%) and no significant publication bias according to the funnel plot (Figure 4), Begg’s test (*z* = 1.68. *P* = 0.093), and Egger’s test (*t* = 1.24, *P* = 0.22).

**Figure 4.**
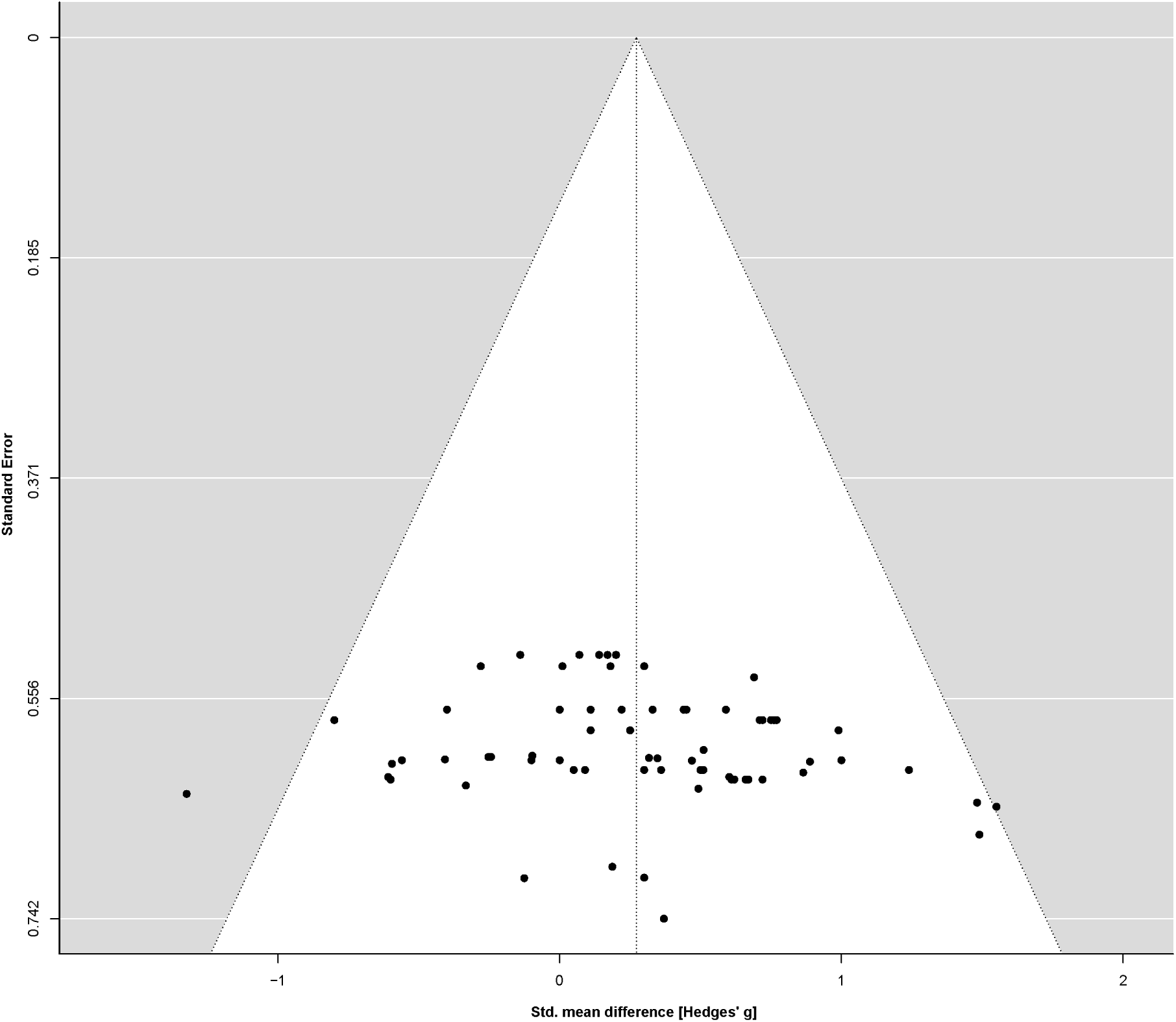
Funnel plot showing the standardized mean differences from all experimental results included in the meta-analysis. The relative symmetry suggests low risk of publication bias.

**Figure 5.**
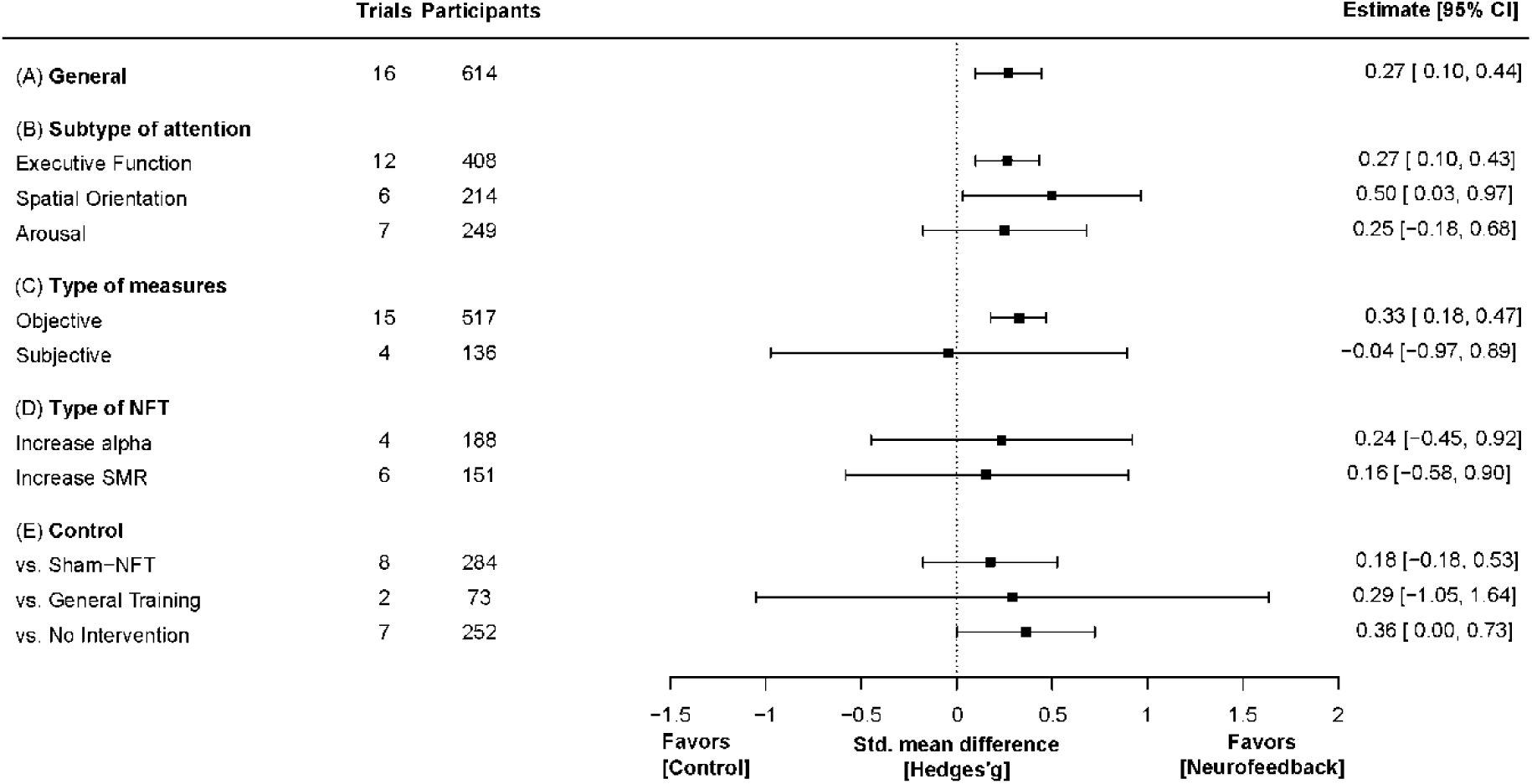
Forest plot of NFT efficacy for improving attentional performance. (A) Overall effect size estimate. (B–E) Estimated effect sizes for each subtype of attentional performance (B), type of outcome measures (C), type of brain activity targeted for NFT (D), and type of control (E). The squares are the means and error bars are the 95% confidence intervals of estimated effect sizes comparing NFT to control conditions.

We performed sensitivity analyses to ensure the robustness of the result. The repeated meta-analysis using a fixed-effects model (instead of the original random-effects model) revealed the small effect size (SMD = 0.27, 95% CI = 0.10–0.44, *t* = 3.77, *P* = 0.0076). The result of the meta-analysis also exhibited the small effect size by including the two studies with high overall risk of bias (18 trials, SMD = 0.25, 95% CI = 0.10–0.41, *t* = 3.80, *P* = 0.0054), by restricting the studies to those performing NFT with EEG (14 trials, SMD = 0.26, 95% CI = 0.03–0.48, *t* = 2.74, *P* = 0.030), or by excluding the effect sizes calculated from the scores after NFT (i.e., not from the change of the score after NFT) (15 trials, SMD = 0.28, 95% CI = 0.10–0.46, *t* = 3.73, *P* = 0.0088).

### 3.6 Meta-regression analyses

Meta-regression analysis revealed no significant relationship between effect size and the total duration of NFT (12 trials; beta = 0.0003, 95% CI = −0.0006–0.0012, t = 1.04, *P* = 0.37; Supplementary Figure 2A) or the number of NFT sessions (15 trials; beta = 0.013, 95% CI = −0.013–0.038, *t* =1.58, *P* = 0.21; Supplementary Figure 2B). We also found no significant correlation between effect size and the publication years (16 trials; beta = 0.0007, 95% CI = −0.035–0.036, *t* =0.05, *P* = 0.96; Supplementary Figure 2C).

### 3.7 Subgroup meta-analyses

We performed subgroup meta-analyses to evaluate the effect of NFT on each subtype of attentional performance (Figure 5B), type of outcome measure (objective or subjective; Figure 5C), type of targeted brain activity to be increased with NFT (Figure 5D), and type of control condition (Figure 5E). The estimated effect on executive function (12 trials) was small (SMD = 0.27, 95% CI = 0.10–0.43, *t* = 4.06, *P* = 0.0088; heterogeneity: Q = 45.8, *P* = 0.72, I^2^ = 0%) and that on spatial orientation (6 trials) was moderate (SMD = 0.50, 95% CI = 0.034–0.97, *t* = 2.96, *P* = 0.041; heterogeneity: Q = 3.12, *P* = 0.87, I^2^ = 0%), while that on arousal (7 trials) was not significant (SMD = 0.25, 95% CI = −0.18–0.68, *t* = 1.58, *P* = 0.18; heterogeneity: Q = 4.99, *P* = 0.96, I^2^ = 0%) (Figure 5B). However, there was no significant difference in NFT effect size among subtypes (*F*_3, 3.34_ = 0.56, *P* = 0.68). Effect size was small regarding the 15 trials using objective measures (SMD = 0.33, 95% CI = 0.19–0.48, *t* = 5.67, *P* = 0.0017; heterogeneity: Q = 42.01, *P* = 0.93, I^2^ = 0%), and not significant across the four trials using subjective measures (SMD = −0.040, 95% CI = −0.97– 0.89, *t* = −0.22, *P* = 0.85; heterogeneity: Q = 6.62, *P* = 0.83, I^2^ = 0%) (Figure 5C). The difference in NFT effect between types of outcome measures was not significant (*F*_1, 2.31_ = 4.16, *P* = 0.16).

When intervention targets were considered separately, NFT demonstrated no significant effect on either alpha power (four trials, SMD = 0.24, 95% CI = −0.45– 0.92, *t* = 1.57, *P* = 0.26; heterogeneity: Q = 16.3, *P* = 0.30, I^2^ = 0%) or SMR power (six trials, SMD = 0.16, 95% CI = −0.58–0.90, *t* = 0.81, *P* = 0.49; heterogeneity: Q = 23.8, *P* = 0.25, I^2^ = 14.7%). We did not evaluate the effect of NFT using other neuroimaging modalities because only one study used NIRS or fMRI.

Notably, NFT had no significant effect on overall attentional performance when compared to sham-NFT as the control condition (eight trials, SMD = 0.18, 95% CI = −0.18–0.53, *t* = 1.57, *P* = 0.21; heterogeneity: Q = 16.9, *P* = 0.98, I^2^ = 0%) (Figure 5E), and also no significant effects on each subtype of attentional performance either (executive function: six trials, SMD = 0.20, 95% CI = −0.16–0.56, *t* = 2.08, *P* = 0.15; spatial orientation: two trials, SMD = 0.09, 95% CI = −2.81–3.00, *t* = 0.39, *P* = 0.76; and arousal: five trials, SMD = 0.22, 95% CI = −0.49–0.92, *t* = 1.05, *P* = 0.38). The effect of NFT was not significantly superior to general training (two trials, SMD = 0.29, 95% CI = −1.05–1.63, *t* = 2.77, *P* = 0.22; heterogeneity: Q = 6.50, *P* = 0.44, I^2^ = 0%). In contrast, NFT was superior to no intervention (seven trials, SMD = 0.36, 95% CI = 0.003–0.73, *t* = 2.90, *P* = 0.049; heterogeneity: Q = 27.5, *P* = 0.44, I^2^ = 0%). Again, there was no significant difference in effect among trials using different control conditions (*F*_2, 1.81_ = 0.43, *P* = 0.70). We found that most subgroup meta-analyses (eight out of 10; other than the studies investigating the effect on executive function and on objective measures) in our study were underpowered (i.e., 1−β was less than 0.8; Supplementary Table 3).

## 4 Discussion

The present study evaluated the effects of NFT on attentional performance using both a qualitative synthesis and meta-analysis of parallel RCTs. We also performed subgroup meta-analyses to assess effects of NFT on specific subsets of attentional performance (executive function, spatial orientation, and arousal) as well as types of outcome measure (subjective or objective), targeted brain activity to be increased with NFT (alpha or SMR power), and control condition (sham-NFT, general training, and no intervention). A search of multiple English and Japanese databases identified 41 studies appropriate for qualitative synthesis and 16 trials (15 studies) acceptable for meta-analyses. The main meta-analysis revealed a significant overall effect of NFT on attentional performance, and subgroup meta-analyses revealed significant effects on executive function and spatial orientation but not on arousal. However, neither the overall effect nor the effects on individual subsets were significant when compared to sham-NFT.

### 4.1 The tendency of the studies revealed from the qualitative synthesis

Since 2017, over half of the studies included in our qualitative synthesis have been published, indicating that research on NFT for improving attentional performance is still growing. This is supported by the fact that most of the studies included here were exploratory, since they did not conduct pre-registration (except for Kim et al., (2019)). However, as the number of publications has increased monotonically over the past five years, more results in this area will be gathered to improve the quality of meta-analysis in the next decade.

Approximately 75% of the studies used EEG to measure brain activity in NFT. This pattern was similar to that found in a recent systematic review of NFT (Onagawa et al., 2023). The reason why EEG-based NFT is more common than fMRI- or NIRS-based NFT can be that EEG is relatively inexpensive compared to fMRI and NIRS. Another reason can be the difference in the length of period since its initial report. To the best of our knowledge, the efficacy of EEG-based NFT was shown approximately 30–40 years (Kamiya, 1968) before that of fMRI-(Yoo and Jolesz, 2002) and NIRS-based NFT (Mihara, 2011). Nevertheless, NIRS is as convenient as EEG to measure brain activity, and fMRI has the advantage of high spatial resolution. Moreover, fMRI-based neurofeedback is gaining attention to examine the causal relationship between brain regions and behavior (Finn, 2021). Thus, as the results of fMRI- or NIRS-based NFT accumulate, it would be tempting to compare the effect on attentional performance across these modalities with a meta-analysis.

The most frequent NFT protocol studied to improve attentional performance was NFT targeted to increase SMR power, followed by that to increase beta, alpha, theta, and gamma power. This pattern differs from that revealed in the recent systematic review of NFT to treat patients with ADHD (Van Doren et al., 2019): The three major protocols for ADHD were NFT to increase SMR power and decrease theta power; to increase beta power and decrease theta power; and to regulate SCP. The most notable difference was the proportion of studies using NFT to regulate SCP (two out of 41 studies in our review, while three out of 10 studies in Van Doren et al., (2019)). This might be because of the difference in the initial NFT protocols reported to increase attentional performance. While, in the studies included in our qualitative synthesis, NFT to increase alpha, SMR, and beta were first demonstrated to increase arousal (Hord et al., 1976), executive function (Vernon et al., 2003), and spatial orientation (Egner and Gruzelier, 2004a), respectively, in the systematic review by Van Doren et al. (2019), NFT to regulate SCP was initially reported to alleviate ADHD-related symptoms. Future research will systematically examine which NFT protocol is best for improving attentional performance.

In this systematic review, we highlighted a major concern regarding NFT studies in this field: several studies provided incomplete information on NFT. For example, of the 41 included studies, six, six, and seven studies did not indicate the intervention period, the number of sessions, and the daily intervention duration, respectively. Moreover, most studies did not report whether any adverse events occurred or not. The detailed NFT protocol is crucial for clarifying the optimal protocol for improving attentional performance, while understanding its possible adverse events is important to evaluate the safety of NFT. Future studies should comply with the recent guidelines for reporting of NFT studies (Ros et al., 2020).

### 4.2 Significant overall NFT effect on attentional performance

Our meta-analysis of 15 studies indicated that NFT was superior to controls for enhancing overall attentional performance but with a small effect size (SMD = 0.27). We confirmed that this result was not severely affected by the choice of the analysis method (random-effects model vs. fixed-effects model) or by the change of the exclusion criteria in the meta-analysis (including the studies with high risk of bias, excluding the effect sizes calculated only from the scores after NFT, or excluding the studies conducting NFT other than EEG). The effect size was comparable to a previous meta-analysis of NFT efficacy for inattention (SMD = 0.38) and hyperactivity-impulsivity (SMD = 0.25) in patients with ADHD compared to controls (Van Doren et al., 2019). In contrast, Da Silva and De Souza reported a larger overall mean effect of NFT on attentional performance compared to our findings (*d’* = 0.61; (Da Silva and De Souza, 2021)). These differences can be explained by the different indices used in these two studies: Hedges’ g used in the current study reflects a relatively small sample size (n = 8–25 in each study), whereas *d’* used in (Da Silva and De Souza, 2021) did not take into account the sample size.

We did not find a significant correlation between the NFT dose and efficacy. Lim et al., (2019) reported that at least 24 sessions are required for NFT to improve attentional performance in patients with ADHD. Considering that only three out of 15 studies included in the meta-analysis conducted NFT for more than 24 sessions and the number of sessions of NFT ranged from 1 to 42 sessions, healthy adults might enhance attentional performance with a rather small number of sessions of NFT compared to patients with ADHD.

### 4.3 Variable NFT effect on each subgroup

The meta-analysis revealed significant effects of NFT on executive function and spatial orientation, consistent with previous systematic reviews showing that NFT can enhance executive function (Da Silva and De Souza, 2021) and visual attention (Ordikhani-Seyedlar et al., 2016). However, in a review by (Viviani and Vallesi, 2021), only 16% of the included studies reported that NFT improved executive function. These discrepancies may be explained by differences in the definition of success: our study focused on improvement of attentional performance measured by behavioral responses, whereas Viviani and Vallesi, (2021) also considered EEG changes compared to its controls. In contrast, we found no significant effect of NFT on arousal compared to controls. We surmise that NFT may not be effective in improving arousal compared to in spatial orientation because the number of studies included in arousal was comparable to that in spatial orientation. Further studies are warranted to address why NFT can improve some attentional domains but not others.

While NFT produced an improvement in attentional performance in general, the meta-analysis revealed no significant effect on individual NFT protocols aimed to increase alpha or SMR power. This is surprising, at least in NFT targeted to increase SMR power, because SMR protocol has long been proposed to enhance attentional performance in healthy adults (Egner and Gruzelier, 2001). These non-significant effects of NFT on alpha and SMR power may be explained by the limited number of studies targeting each band power (four trials for alpha and six for SMR power). Another potential reason is that enhancement of a given band power may not necessarily improve all subsets of attentional performance consistently. For example, several studies have suggested that NFT protocols enhancing low beta and SMR power influence distinct aspects of attentional performance (Egner and Gruzelier, 2004a, 2004b).

The meta-analysis also revealed no significant effect of NFT on attentional performance when compared specifically to sham-NFT or to general training. Arina and colleagues (2017) reported that even sham-NFT can induce unspecific effect such as reducing tension or anxiety. This finding suggests that NFT effects specific to modulation of certain EEG band power cannot be evaluated only by comparison with a non-intervention group. Nevertheless, only about half of the studies (21 out of 41) compared NFT effect with sham-NFT, and the details of the blinding protocols were unclear in most studies. As suggested in other systematic reviews on NFT (Da Silva and De Souza, 2021; Rogala et al., 2016), stringent triple-blind parallel RCTs are needed to compare the effects of NFT with sham-NFT.

### 4.4 Limitations

The present study has a potential limitation: we cannot assess the NFT effect on each neuroimaging modality. This is because only one study conducted NFT with NIRS (Nouchi et al., 2021) or fMRI (Kim et al., 2019) to enhance attentional performance. NIRS is a convenient tool to measure brain activity as EEG, and several studies suggest that NFT with NIRS is useful for the rehabilitation after stroke (Mihara et al., 2021; Rahman et al., 2020). NFT with fMRI is also a promising tool to modulate brain activity (Koizumi et al., 2020). Future studies are required to determine which neuroimaging modality can best improve attentional performance.

Another limitation of this study was that it was not pre-registered. Therefore, we performed meta-regression analysis and subgroup meta-analysis exploratory. As this might introduce publication bias, these results should be interpreted with caution. We also acknowledge the limited number of trials in our subgroup meta-analyses. If the number of trials included in a meta-analysis is small (e.g., ≤ five trials (Zhou and Shen, 2022)), the result of that analysis must be cautiously interpreted. Further studies will be required to validate which subtype of attentional performance is more benefitted by NFT, which NFT protocol can most effectively improve attentional performance, and whether NFT is beneficial compared to sham-NFT or general training.

Aside from these limitations, it should be noted that only eight trials were included in the meta-analysis that compared the effect of NFT with that of sham-NFT. This is because the effect of the studies that compared with no intervention can include placebo effects, indicating that the effect of NFT in these trials could be influenced by the participant’s motivation or expectation towards neurofeedback (Thibault et al., 2017). In fact, one study reported that the subjective sleep quality scores improved with sham-NFT, even though sham-NFT did not regulate neural activity (Schabus et al., 2017; Thibault et al., 2017). Therefore, a comparison with sham-NFT will be essential to reveal the effects specific to the modulation of the targeted brain activity.

Another thing to note is that only two trials compared NFT effect with general training. Comparison with general training is also crucial to clarify which training (NFT or other) is more useful for enhancing attentional performance. Taken together, more studies comparing the NFT effect with sham-NFT and general training are needed to scientifically validate the NFT effect on attentional performance and the superiority of NFT over alternative methods.

### 4.5 Conclusion

From the currently available studies, NFT significantly enhances attentional performance compared to its control, without a significant dose–response relationship in healthy adults. Furthermore, NFT has been observed to significantly enhance executive function and spatial orientation, although whether NFT can significantly enhance arousal is still unknown. Unfortunately, approximately half of the studies included in our meta-analysis did not compare NFT to sham-NFT, and the superiority of NFT over sham-NFT and alternative methods for enhancing attentional performance remains uncertain. To address these gaps, future multicenter parallel RCTs with large sample sizes are required to compare NFT effect with sham-NFT in a rigorous triple-blinded design, as well as with non-NFT training.

## Supporting information

https://www.biorxiv.org/content/biorxiv/early/2023/03/22/2023.03.19.533384/DC1/embed/media-1.pdf?download=true

## Data and code availability

The data (mean(s) and standard deviation(s) for experimental and control groups and the converted effect size(s) for each study) and codes used in this study were shared in OSF (https://osf.io/5ze7v/).

## Author contributions

Ikko Kimura: Methodology, Investigation, Data Curation, Formal analysis, Writing – Original Draft, Visualization

Hiroki Noyama: Methodology, Investigation, Data Curation, Formal analysis, Writing – Original Draft, Visualization

Ryoji Onagawa: Investigation, Data Curation, Writing – Review & Editing

Mitsuaki Takemi: Conceptualization, Project Administration, Writing – Review & Editing

Rieko Osu: Supervision, Writing – Review & Editing

Jun-ichiro Kawahara: Supervision, Project Administration, Writing – Review & Editing

## Acknowledgements

This work was conducted as a part of Braintech Guidebook development in JST Moonshot R&D to MT (Grant Number JPMJMS2012). We thank the members of the Evidence Evaluation Committee for Braintech Guidebook, a specially organized group for the Moonshot R&D project, for their comments on the manuscript.

