## Supplementary material for "Efficacy of neurofeedback training for improving attentional performance in healthy adults: A systematic review and meta-analysis": https://www.biorxiv.org/content/biorxiv/early/2023/03/22/2023.03.19.533384/DC1/embed/media-1.pdf?download=true: KimuraI2023ImagingNeuroscience_Suppl_20230904_MERGED.pdf

**Supplementary Table 1.** Search query for each database.

| Database | Formula |
|---|---|
| PubMed | #1. Attention[Mesh] OR "Executive Function"[Mesh] OR Wakefulness[Mesh]<br>#2. "Biofeedback, Psychology"[Mesh] OR "Brain-Computer Interfaces"[Mesh]<br>#3. english[Language]<br>#4. "Mental Disorders"[Mesh] OR Rehabilitation[Mesh] OR "Brain Diseases"[Mesh]<br>#5. animal*[TI] OR rodent*[TI] OR monkey*[TI] OR rat[TI] OR rats[TI] OR macaque*[TI]<br>#6. #1 AND #2 AND #3 NOT #4 NOT #5 |
| Scopus | ( TITLE-ABS-KEY ( attention* OR wakefulness OR {executive function} ) ) AND ( TITLE-ABS-KEY ( neurofeedback OR {neuro feedback} OR "brain machine interface" OR "brain computer interface" OR ( feedback AND ( eeg OR electroencephalogra* ) ) ) ) AND NOT ( TITLE ( adhd OR disorder* OR deficit* OR patient* OR rehabilitation OR epilep* OR disease* OR depress* OR injur* OR damage* ) ) AND NOT ( TITLE ( animal* OR rodent* OR monkey* OR rat OR rats OR macaque* ) ) AND ( LIMIT-TO ( DOCTYPE , "ar" ) OR LIMIT-TO ( DOCTYPE , "cp" ) ) AND ( LIMIT-TO ( LANGUAGE , "English" ) ) AND ( LIMIT-TO ( SRCTYPE , "j" ) OR LIMIT-TO ( SRCTYPE , "p" ) ) |
| Web of science | #1. TS=(attention* OR wakefulness OR "executive function")<br>#2. TS=(neurofeedback OR "neuro feedback" OR "brain machine interface" OR "brain computer interface" OR (feedback AND (EEG OR electroencephalogra*)))<br>#3. LA=(English)<br>#4. TI=(adhd OR disorder* OR deficit* OR patient* OR rehabilitation OR epilep* OR disease* OR depress* OR injur* OR damage*)<br>#5. TI=(animal* OR rodent* OR monkey* OR rat OR rats OR macaque*)<br>#6. #1 AND #2 AND #3 NOT #4 NOT #5 (Document Type=(article OR letter OR "proceedings paper")) |
| APA PsycInfo | #1. (attention* OR wakefulness OR "executive function") in Any Field<br>#2. (neurofeedback OR "neuro feedback" OR "brain machine interface" OR "brain computer interface" OR (feedback AND (EEG OR electroencephalogra*))) in Any Field<br>#3. (adhd OR disorder* OR deficit* OR patient* OR rehabilitation OR epilep* OR disease* OR depress* OR injur* OR damage*) in Title<br>#4. (animal* OR rodent* OR monkey* OR rat OR rats OR macaque*) in Title<br>#5. #1 AND #2 NOT #3 NOT #4<br>#6. Age Group: Young Adulthood (18-29 yrs) OR Thirties (30-39 yrs) OR Middle Age (40-64 yrs)<br>#7. Language: English |

| | |
|---|---|
| | #8. Publication Type: Peer-reviewed<br>#9. #5 AND #6 AND #7 AND #8 |
| JDreamIII | L1 "注意"/AL OR "アテンション"/AL OR "注意力"/AL OR "注意集中"/AL OR "注目"/AL OR "注視"/AL OR "注視行動"/AL OR "用心"/AL OR "警戒"/AL OR "覚醒"/AL OR "実行機能"/AL OR "遂行機能"/AL<br>L2 (フィードバック AND (eeg + 脳波)) +ニューロフィードバック+ブレインマシンインターフェース +ブレインコンピュータインターフェース<br>L3 精神疾患 OR "精神病"/AL OR "こころの病気"/AL OR "精神機能障害"/AL OR "精神疾患"/AL OR "精神病性障害"/AL OR "精神病状態"/AL<br>L4 子供+患者+子ども+高齢者<br>L4 (L1 AND L2 AND (JA/LA) AND (a1/DT) NOT L3 NOT L4) |
| Ichu-shi | #1. [注意(心理学)]/TH OR [実行機能(意識過程)]/TH OR [覚醒状態]/TH<br>#2. バイオフィードバック/TH or ニューロフィードバック/TH or [ブレイン－マシンインターフェース]/TH<br>#3. 精神疾患/TH OR [脳疾患]/TH<br>#4. (#1 and #2 not #3) and (LA=日本語 PT=原著論文 CK=成人(19～44),中年(45～64)) |

**Supplementary Table 2.** The criteria for each domain of risk of bias

| | Low | High |
|---|---|---|
| D1: Random sequence generation (selection bias) | The random allocation is based on a random number table, coin toss, or drawing. | The allocation is based on birth date, date of visit, or ID of the participant, or the judgment of the allocation includes the experimenters' or participants' thoughts. |
| D2: Allocation concealment (selection bias) | Participants and researchers cannot predict allocation. | Participants and researchers can predict allocation. |
| D3: Blinding of participants and personnel (performance bias) | The blinding of participants and experimenters who do neurofeedback is assured. | The blinding of participants or experimenters who do neurofeedback is not done or incomplete. |
| D4: Blinding outcome assessment (detection bias) | The blinding of the evaluator of the outcome is assured. | The blinding of the evaluator of the outcome is not done or incomplete. |
| D5: Incomplete outcome data (attrition bias) | The data from all participants were obtained or the proportion of missing data is small ( < 10% for each group). The reasons for missing data are unlikely to affect the estimates of the effect on the intervention. | The proportion of missing data is high. The reasons for missing data are likely to affect the estimates of the effect on the intervention. The proportions or reasons for missing data are different between neurofeedback and control groups. |
| D6: Selective reporting (reporting bias) | All of the primary and secondary outcomes were reported as described in the pre-registration. | Not all of the outcomes were reported as described in the pre-registration. |
| D7: Other bias | No other possible causes of bias. | Has potential cause of bias, such as early termination of the study, conflict of interest, and inadequate statistical methods. |

**Supplementary Table 3.** Summary of the power analyses. The estimated power and required number of sample sizes were estimated from the standard mean difference (SMD) and tau^2 estimated from each meta-analysis.

| **Group conditions** | **SMD** | $\tau^2$ | **Estimated power** | **Required number of sample sizes [number of RCTs (number of participants)]** |
|---|---|---|---|---|
| General | 0.27 | 0.00 | 0.88 | 13 (442) |
| Subtype: Executive function | 0.27 | 0.00 | 0.77 | 13 (442) |
| Subtype: Spatial orientation | 0.50 | 0.00 | 0.94 | 4 (136) |
| Subtype: Arousal | 0.25 | 0.00 | 0.48 | 16 (544) |
| Measures: Objective | 0.33 | 0.00 | 0.96 | 9 (306) |
| Measures: Subjective | -0.05 | 0.00 | 0.056 | NA (NA) |
| Intervention: Alpha | 0.24 | 0.00 | 0.28 | 17 (578) |
| Intervention: SMR | 0.16 | 0.067 | 0.19 | 40 (1360) |
| Comparison: Sham-NFT | 0.18 | 0.00 | 0.31 | 30 (1020) |
| Comparison: General training | 0.29 | 0.00 | 0.22 | 12 (408) |
| Comparison: No intervention | 0.36 | 0.0021 | 0.78 | 8 (272) |

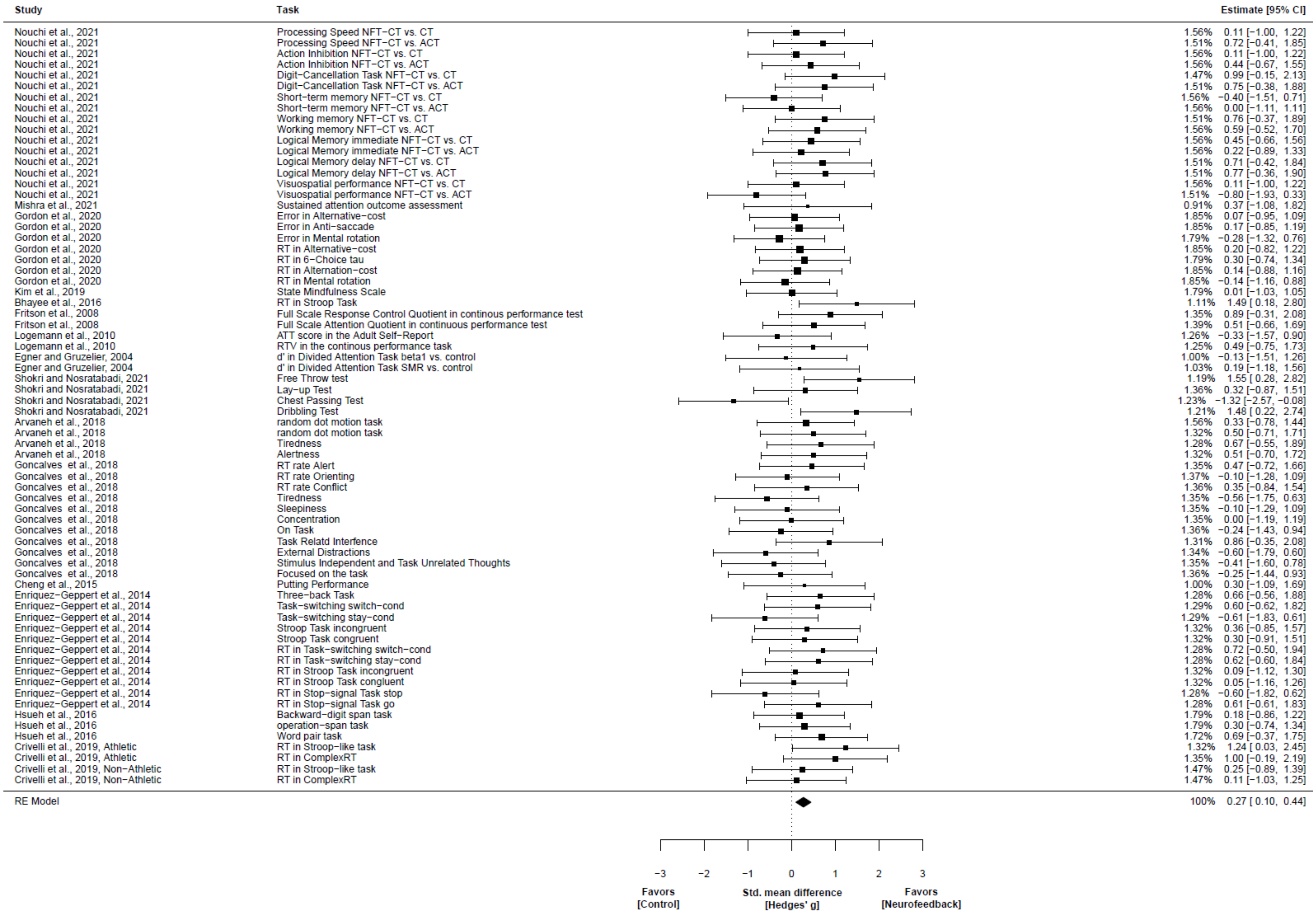
Study
Task
Estimate [95% CI]
Nouchi et al., 2021 Processing Speed NFT−CT vs. CT 1.56% 0.11 [−1.00, 1.22]
Nouchi et al., 2021 Processing Speed NFT−CT vs. ACT 1.51% 0.72 [−0.41, 1.85]
Nouchi et al., 2021 Action Inhibition NFT−CT vs. CT 1.56% 0.11 [−1.00, 1.22]
Nouchi et al., 2021 Action Inhibition NFT−CT vs. ACT 1.56% 0.44 [−0.67, 1.55]
Nouchi et al., 2021 Digit−Cancellation Task NFT−CT vs. CT 1.47% 0.99 [−0.15, 2.13]
Nouchi et al., 2021 Digit−Cancellation Task NFT−CT vs. ACT 1.51% 0.75 [−0.38, 1.88]
Nouchi et al., 2021 Short−term memory NFT−CT vs. CT 1.56% −0.40 [−1.51, 0.71]
Nouchi et al., 2021 Short−term memory NFT−CT vs. ACT 1.56% 0.00 [−1.11, 1.11]
Nouchi et al., 2021 Working memory NFT−CT vs. CT 1.51% 0.76 [−0.37, 1.89]
Nouchi et al., 2021 Working memory NFT−CT vs. ACT 1.56% 0.59 [−0.52, 1.70]
Nouchi et al., 2021 Logical Memory immediate NFT−CT vs. CT 1.56% 0.45 [−0.66, 1.56]
Nouchi et al., 2021 Logical Memory immediate NFT−CT vs. ACT 1.56% 0.22 [−0.89, 1.33]
Nouchi et al., 2021 Logical Memory delay NFT−CT vs. CT 1.51% 0.71 [−0.42, 1.84]
Nouchi et al., 2021 Logical Memory delay NFT−CT vs. ACT 1.51% 0.77 [−0.36, 1.90]
Nouchi et al., 2021 Visuospatial performance NFT−CT vs. CT 1.56% 0.11 [−1.00, 1.22]
Nouchi et al., 2021 Visuospatial performance NFT−CT vs. ACT 1.51% −0.80 [−1.93, 0.33]
Mishra et al., 2021 Sustained attention outcome assessment 0.91% 0.37 [−1.08, 1.82]
Gordon et al., 2020 Error in Alternative−cost 1.85% 0.07 [−0.95, 1.09]
Gordon et al., 2020 Error in Anti−saccade 1.85% 0.17 [−0.85, 1.19]
Gordon et al., 2020 Error in Mental rotation 1.79% −0.28 [−1.32, 0.76]
Gordon et al., 2020 RT in Alternative−cost 1.85% 0.20 [−0.82, 1.22]
Gordon et al., 2020 RT in 6−Choice tau 1.79% 0.30 [−0.74, 1.34]
Gordon et al., 2020 RT in Alternation−cost 1.85% 0.14 [−0.88, 1.16]
Gordon et al., 2020 RT in Mental rotation 1.85% −0.14 [−1.16, 0.88]
Kim et al., 2019 State Mindfulness Scale 1.79% 0.01 [−1.03, 1.05]
Bhayee et al., 2016 RT in Stroop Task 1.11% 1.49 [0.18, 2.80]
Fritson et al., 2008 Full Scale Response Control Quotient in continous performance test 1.35% 0.89 [−0.31, 2.08]
Fritson et al., 2008 Full Scale Attention Quotient in continuous performance test 1.39% 0.51 [−0.66, 1.69]
Logemann et al., 2010 ATT score in the Adult Self−Report 1.26% −0.33 [−1.57, 0.90]
Logemann et al., 2010 RTV in the continous performance task 1.25% 0.49 [−0.75, 1.73]
Egner and Gruzelier, 2004 d' in Divided Attention Task beta1 vs. control 1.00% −0.13 [−1.51, 1.26]
Egner and Gruzelier, 2004 d' in Divided Attention Task SMR vs. control 1.03% 0.19 [−1.18, 1.56]
Shokri and Nosratabadi, 2021 Free Throw test 1.19% 1.55 [0.28, 2.82]
Shokri and Nosratabadi, 2021 Lay−up Test 1.36% 0.32 [−0.87, 1.51]
Shokri and Nosratabadi, 2021 Chest Passing Test 1.23% −1.32 [−2.57, −0.08]
Shokri and Nosratabadi, 2021 Dribbling Test 1.21% 1.48 [0.22, 2.74]
Arvaneh et al., 2018 random dot motion task 1.56% 0.33 [−0.78, 1.44]
Arvaneh et al., 2018 random dot motion task 1.32% 0.50 [−0.71, 1.71]
Arvaneh et al., 2018 Tiredness 1.28% 0.67 [−0.55, 1.89]
Arvaneh et al., 2018 Alertness 1.32% 0.51 [−0.70, 1.72]
Goncalves et al., 2018 RT rate Alert 1.35% 0.47 [−0.72, 1.66]
Goncalves et al., 2018 RT rate Orienting 1.37% −0.10 [−1.28, 1.09]
Goncalves et al., 2018 RT rate Conflict 1.36% 0.35 [−0.84, 1.54]
Goncalves et al., 2018 Tiredness 1.35% −0.56 [−1.75, 0.63]
Goncalves et al., 2018 Sleepiness 1.35% −0.10 [−1.29, 1.09]
Goncalves et al., 2018 Concentration 1.35% 0.00 [−1.19, 1.19]
Goncalves et al., 2018 On Task 1.36% −0.24 [−1.43, 0.94]
Goncalves et al., 2018 Task Relatd Interfence 1.31% 0.86 [−0.35, 2.08]
Goncalves et al., 2018 External Distractions 1.34% −0.60 [−1.79, 0.60]
Goncalves et al., 2018 Stimulus Independent and Task Unrelated Thoughts 1.35% −0.41 [−1.60, 0.78]
Goncalves et al., 2018 Focused on the task 1.36% −0.25 [−1.44, 0.93]
Cheng et al., 2015 Putting Performance 1.00% 0.30 [−1.09, 1.69]
Enriquez−Geppert et al., 2014 Three−back Task 1.28% 0.66 [−0.56, 1.88]
Enriquez−Geppert et al., 2014 Task−switching switch−cond 1.29% 0.60 [−0.62, 1.82]
Enriquez−Geppert et al., 2014 Task−switching stay−cond 1.29% −0.61 [−1.83, 0.61]
Enriquez−Geppert et al., 2014 Stroop Task incongruent 1.32% 0.36 [−0.85, 1.57]
Enriquez−Geppert et al., 2014 Stroop Task congruent 1.32% 0.30 [−0.91, 1.51]
Enriquez−Geppert et al., 2014 RT in Task−switching switch−cond 1.28% 0.72 [−0.50, 1.94]
Enriquez−Geppert et al., 2014 RT in Task−switching stay−cond 1.28% 0.62 [−0.60, 1.84]
Enriquez−Geppert et al., 2014 RT in Stroop Task incongruent 1.32% 0.09 [−1.12, 1.30]
Enriquez−Geppert et al., 2014 RT in Stroop Task congluent 1.32% 0.05 [−1.16, 1.26]
Enriquez−Geppert et al., 2014 RT in Stop−signal Task stop 1.28% −0.60 [−1.82, 0.62]
Enriquez−Geppert et al., 2014 RT in Stop−signal Task go 1.28% 0.61 [−0.61, 1.83]
Hsueh et al., 2016 Backward−digit span task 1.79% 0.18 [−0.86, 1.22]
Hsueh et al., 2016 operation−span task 1.79% 0.30 [−0.74, 1.34]
Hsueh et al., 2016 Word pair task 1.72% 0.69 [−0.37, 1.75]
Crivelli et al., 2019, Athletic RT in Stroop−like task 1.32% 1.24 [0.03, 2.45]
Crivelli et al., 2019, Athletic RT in ComplexRT 1.35% 1.00 [−0.19, 2.19]
Crivelli et al., 2019, Non−Athletic RT in Stroop−like task 1.47% 0.25 [−0.89, 1.39]
Crivelli et al., 2019, Non−Athletic RT in ComplexRT 1.47% 0.11 [−1.03, 1.25]
RE Model 100% 0.27 [0.10, 0.44]
−3 −2 −1 0 1 2 3
Favors [Control]
Std. mean difference [Hedges' g]
Favors [Neurofeedback]

**Supplementary Figure 1.** Forest plot regarding the NFT effect in improving attentional performance. The errorbar shows the estimated standardized mean difference (Hedges' g) and 95% confidence interval of each outcome included in the meta-analysis.

Abbreviations: NFT, neuro-feedback training; CT, cognitive training; ACT, active control; RT, reaction time; RTV, reaction time variability; ATT score, scores for attentional problems

A

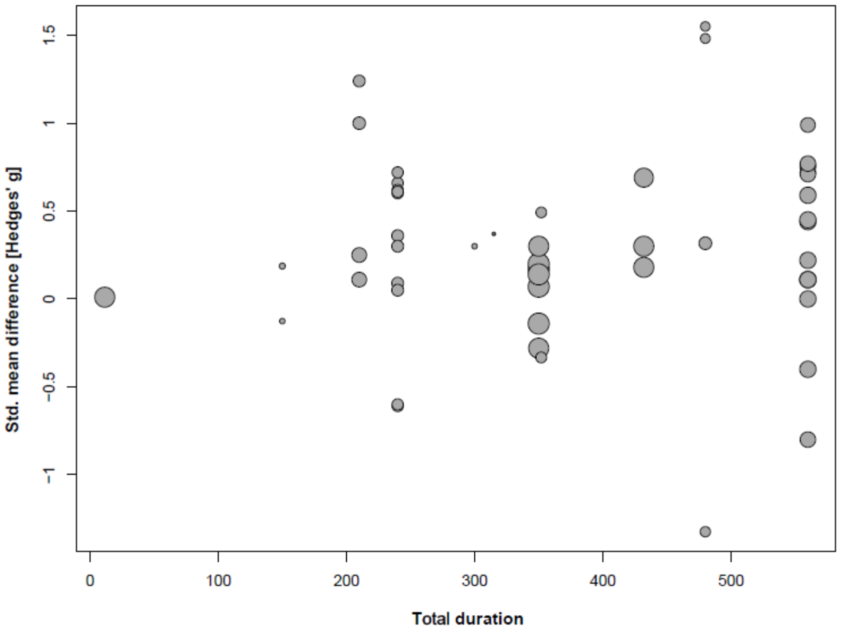

Std. mean difference [Hedges' g]
1.5
1
0.5
0
-0.5
-1
0
100
200
300
400
500
Total duration


B

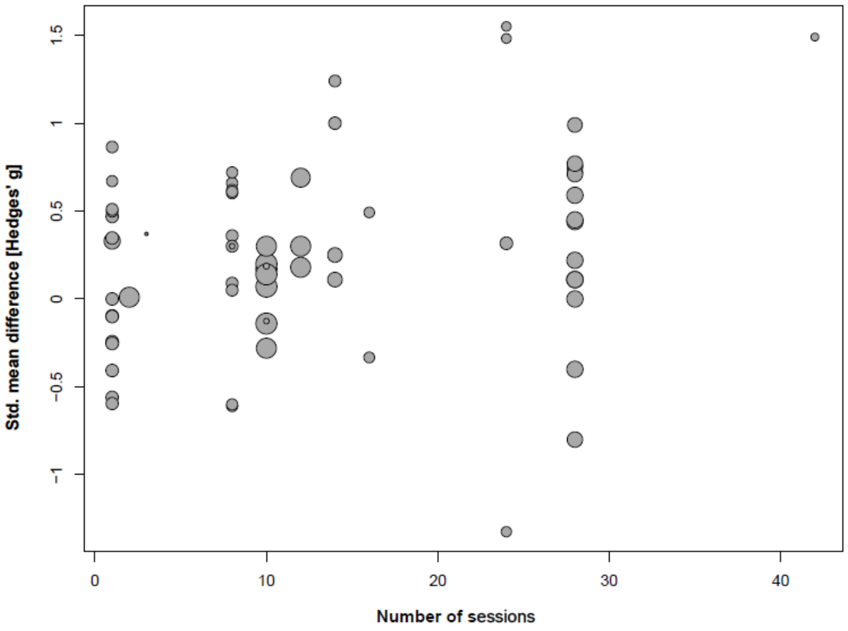

Std. mean difference [Hedges' g]
1.5
1
0.5
0
-0.5
-1
0
10
20
30
40
Number of sessions


C

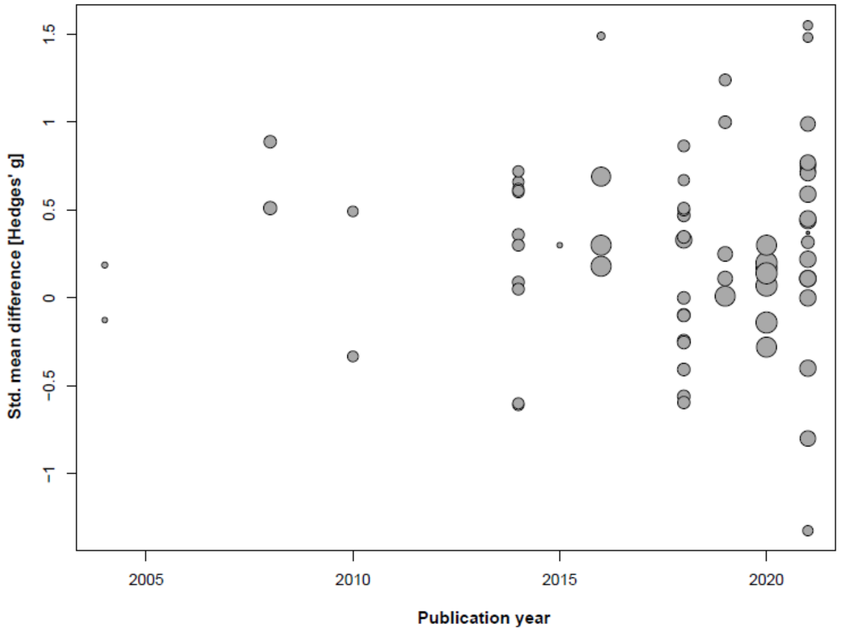

Std. mean difference [Hedges' g]
1.5
1
0.5
0
-0.5
-1
2005
2010
2015
2020
Publication year

**Supplementary Figure 2.** Relationship between the effects on attentional performance and (A) the total duration of NFT, (B) the number of NFT sessions, and (C) the publication year. Each dot shows the results from each trial.

### PRISMA 2020 for Abstracts Checklist

| Section and Topic | Item # | Checklist item | Reported (Yes/No) |
|---|---|---|---|
| **TITLE** | | | |
| Title | 1 | Identify the report as a systematic review. | Yes |
| **BACKGROUND** | | | |
| Objectives | 2 | Provide an explicit statement of the main objective(s) or question(s) the review addresses. | Yes |
| **METHODS** | | | |
| Eligibility criteria | 3 | Specify the inclusion and exclusion criteria for the review. | No |
| Information sources | 4 | Specify the information sources (e.g. databases, registers) used to identify studies and the date when each was last searched. | Yes |
| Risk of bias | 5 | Specify the methods used to assess risk of bias in the included studies. | Yes |
| Synthesis of results | 6 | Specify the methods used to present and synthesise results. | Yes |
| **RESULTS** | | | |
| Included studies | 7 | Give the total number of included studies and participants and summarise relevant characteristics of studies. | Yes |
| Synthesis of results | 8 | Present results for main outcomes, preferably indicating the number of included studies and participants for each. If meta-analysis was done, report the summary estimate and confidence/credible interval. If comparing groups, indicate the direction of the effect (i.e. which group is favoured). | Yes |
| **DISCUSSION** | | | |
| Limitations of evidence | 9 | Provide a brief summary of the limitations of the evidence included in the review (e.g. study risk of bias, inconsistency and imprecision). | Yes |
| Interpretation | 10 | Provide a general interpretation of the results and important implications. | Yes |
| **OTHER** | | | |
| Funding | 11 | Specify the primary source of funding for the review. | No |
| Registration | 12 | Provide the register name and registration number. | No |

*From:* Page MJ, McKenzie JE, Bossuyt PM, Boutron I, Hoffmann TC, Mulrow CD, et al. The PRISMA 2020 statement: an updated guideline for reporting systematic reviews. BMJ 2021;372:n71. doi: 10.1136/bmj.n71

For more information, visit: http://www.prisma-statement.org/

### PRISMA 2020 Checklist

| Section and Topic | Item # | Checklist item | Location where item is reported |
|---|---|---|---|
| **TITLE** | | | |
| Title | 1 | Identify the report as a systematic review. | Title page |
| **ABSTRACT** | | | |
| Abstract | 2 | See the PRISMA 2020 for Abstracts checklist. | Abstract |
| **INTRODUCTION** | | | |
| Rationale | 3 | Describe the rationale for the review in the context of existing knowledge. | Section 1 |
| Objectives | 4 | Provide an explicit statement of the objective(s) or question(s) the review addresses. | Section 1 |
| **METHODS** | | | |
| Eligibility criteria | 5 | Specify the inclusion and exclusion criteria for the review and how studies were grouped for the syntheses. | Section 2.2 and 2.3 |
| Information sources | 6 | Specify all databases, registers, websites, organisations, reference lists and other sources searched or consulted to identify studies. Specify the date when each source was last searched or consulted. | Section 2.1 |
| Search strategy | 7 | Present the full search strategies for all databases, registers and websites, including any filters and limits used. | Section 2.1 and Supp. |
| Selection process | 8 | Specify the methods used to decide whether a study met the inclusion criteria of the review, including how many reviewers screened each record and each report retrieved, whether they worked independently, and if applicable, details of automation tools used in the process. | Section 2.2 |
| Data collection process | 9 | Specify the methods used to collect data from reports, including how many reviewers collected data from each report, whether they worked independently, any processes for obtaining or confirming data from study investigators, and if applicable, details of automation tools used in the process. | Section 2.3 |
| Data items | 10a | List and define all outcomes for which data were sought. Specify whether all results that were compatible with each outcome domain in each study were sought (e.g. for all measures, time points, analyses), and if not, the methods used to decide which results to collect. | Section 2.3 |
| | 10b | List and define all other variables for which data were sought (e.g. participant and intervention characteristics, funding sources). Describe any assumptions made about any missing or unclear information. | Section 2.3 and 2.4 |
| Study risk of bias assessment | 11 | Specify the methods used to assess risk of bias in the included studies, including details of the tool(s) used, how many reviewers assessed each study and whether they worked independently, and if applicable, details of automation tools used in the process. | Section 2.4 |
| Effect measures | 12 | Specify for each outcome the effect measure(s) (e.g. risk ratio, mean difference) used in the synthesis or presentation of results. | Section 2.3 |
| Synthesis methods | 13a | Describe the processes used to decide which studies were eligible for each synthesis (e.g. tabulating the study intervention characteristics and comparing against the planned groups for each synthesis (item #5)). | Section 2.2 |
| | 13b | Describe any methods required to prepare the data for presentation or synthesis, such as handling of missing summary statistics, or data conversions. | Section 2.3 |
| | 13c | Describe any methods used to tabulate or visually display results of individual studies and syntheses. | Section 2.5 |
| | 13d | Describe any methods used to synthesize results and provide a rationale for the choice(s). If meta-analysis was performed, describe the model(s), method(s) to identify the presence and extent of statistical heterogeneity, and software package(s) used. | Section 2.5 |
| | 13e | Describe any methods used to explore possible causes of heterogeneity among study results (e.g. subgroup analysis, meta-regression). | Section 2.5 |
| | 13f | Describe any sensitivity analyses conducted to assess robustness of the synthesized results. | Section 2.5 |
| Reporting bias assessment | 14 | Describe any methods used to assess risk of bias due to missing results in a synthesis (arising from reporting biases). | Section 2.4 |

#### PRISMA 2020 Checklist

| Section and Topic | Item # | Checklist item | Location where item is reported |
|---|---|---|---|
| Certainty assessment | 15 | Describe any methods used to assess certainty (or confidence) in the body of evidence for an outcome. | Section 2.5 |
| **RESULTS** | | | |
| Study selection | 16a | Describe the results of the search and selection process, from the number of records identified in the search to the number of studies included in the review, ideally using a flow diagram. | Section 3.1 |
| | 16b | Cite studies that might appear to meet the inclusion criteria, but which were excluded, and explain why they were excluded. | Section 3.1 |
| Study characteristics | 17 | Cite each included study and present its characteristics. | Table |
| Risk of bias in studies | 18 | Present assessments of risk of bias for each included study. | Section 3.4 |
| Results of individual studies | 19 | For all outcomes, present, for each study: (a) summary statistics for each group (where appropriate) and (b) an effect estimate and its precision (e.g. confidence/credible interval), ideally using structured tables or plots. | Section 3.5, 3.6, and Supp. |
| Results of syntheses | 20a | For each synthesis, briefly summarise the characteristics and risk of bias among contributing studies. | Section 3.4 |
| | 20b | Present results of all statistical syntheses conducted. If meta-analysis was done, present for each the summary estimate and its precision (e.g. confidence/credible interval) and measures of statistical heterogeneity. If comparing groups, describe the direction of the effect. | Section 3.2, 3.5, and 3.6 |
| | 20c | Present results of all investigations of possible causes of heterogeneity among study results. | Section 3.5 and 3.6 |
| | 20d | Present results of all sensitivity analyses conducted to assess the robustness of the synthesized results. | Section 3.5 |
| Reporting biases | 21 | Present assessments of risk of bias due to missing results (arising from reporting biases) for each synthesis assessed. | Section 3.4 |
| Certainty of evidence | 22 | Present assessments of certainty (or confidence) in the body of evidence for each outcome assessed. | Section 3.5 |
| **DISCUSSION** | | | |
| Discussion | 23a | Provide a general interpretation of the results in the context of other evidence. | Section 4.1 and 4.2 |
| | 23b | Discuss any limitations of the evidence included in the review. | Section 4.3 |
| | 23c | Discuss any limitations of the review processes used. | Section 4.3 |
| | 23d | Discuss implications of the results for practice, policy, and future research. | Section 4.4 |
| **OTHER INFORMATION** | | | |
| Registration and protocol | 24a | Provide registration information for the review, including register name and registration number, or state that the review was not registered. | Section 2 |
| | 24b | Indicate where the review protocol can be accessed, or state that a protocol was not prepared. | Availability statement |
| | 24c | Describe and explain any amendments to information provided at registration or in the protocol. | N.A. |
| Support | 25 | Describe sources of financial or non-financial support for the review, and the role of the funders or sponsors in the review. | Acknowledgement |
| Competing interests | 26 | Declare any competing interests of review authors. | Competing interests |
| Availability of data, code and other | 27 | Report which of the following are publicly available and where they can be found: template data collection forms; data extracted from included studies; data used for all analyses; analytic code; any other materials used in the review. | Availability statemen |

PRISMA 2020 Checklist

| Section and Topic | Item # | Checklist item | Location where item is reported |
|---|---|---|---|
| materials | | | |

*From:* Page MJ, McKenzie JE, Bossuyt PM, Boutron I, Hoffmann TC, Mulrow CD, et al. The PRISMA 2020 statement: an updated guideline for reporting systematic reviews. BMJ 2021;372:n71. doi: 10.1136/bmj.n71

For more information, visit: http://www.prisma-statement.org/
